# Dominance between self-incompatibility alleles determines the mating system of Capsella allopolyploids

**DOI:** 10.1101/2023.04.17.537155

**Authors:** Tianlin Duan, Zebin Zhang, Mathieu Genete, Céline Poux, Adrien Sicard, Martin Lascoux, Vincent Castric, Xavier Vekemans

**Affiliations:** Department of Ecology and Genetics, Evolutionary Biology Centre and Science for Life Laboratory, Uppsala University, S-75236 Uppsala, Sweden; University of Lille, CNRS, UMR 8198 – Evo-Eco-Paleo, F-59000 Lille, France; Department of Plant Biology, Swedish University of Agricultural Sciences, S-750 07 Uppsala, Sweden

**Keywords:** Self-incompatibility, polyploidy, S-locus, *SRK*, *SCR*, genetic dominance, Capsella

## Abstract

The shift from outcrossing to self-fertilization is one of the main evolutionary transitions in plants, and has broad effects on evolutionary trajectories. In Brassicaceae, the ability to impede self-fertilization is controlled by two genes, *SCR* and *SRK,* tightly linked within the S-locus. A series of small non-coding RNAs also encoded within the S-locus regulates the transcriptional activity of *SCR* alleles, resulting in a linear dominance hierarchy between them. Brassicaceae allopolyploid species are often self-compatible (SC) even when one of their parents is self-incompatible, but the causes of the loss of self-incompatibility (SI) in polyploid lineages have generally remained elusive. We used a series of synthetic hybrids obtained between self-fertilizing *Capsella orientalis* and outcrossing *C. grandiflora* to test whether the breakdown of SI in allopolyploid species, such as *C. bursa-pastoris*, could be explained by the dominance interactions between S-haplotypes inherited from the parental lineages. After establishing a database of reference S-allele sequences, we used RNA-sequencing data from young inflorescences to measure allele-specific expression of the *SCR* and *SRK* genes in diploid and tetraploid synthetic hybrids. We then compared the observed expression of *SCR* alleles with the predicted dominance relationship between S-haplotypes in pollen and with the seed set from autonomous self-fertilization in the synthetic hybrids. Our results formally establish that upon hybridization, the immediate effect on the mating system depends on the relative dominance between S-alleles inherited from the parental species. They illustrate that a detailed understanding of the genetic architecture of the control of SI is essential to predict the patterns of association between the mating system and changes in ploidy.

**Lay summary:** Polyploidy is the inheritable condition of carrying more than two sets of chromosomes. It can result from within-species genome duplication (auto-polyploidy), or from the merging of sets of chromosomes from different species following hybridization (allo-polyploidy). Because sexual reproduction between individuals of different levels of ploidy is generally not successful, self-fertilization has been considered a key component of the establishment success of polyploid lineages. However, the reasons why the mating system of polyploids may differ from that of their parental species remains mysterious. In plants of the Brassicaceae family, several allopolyploid species arose from hybridization between an outcrossing and a self-fertilizing species, and in most cases the resulting lineages are self-fertilizing. It has been proposed that the mating system of these allopolyploids depends on the dominance relationships between the functional and non-functional self-incompatibility alleles inherited from the parental species. Here, we tested this prediction by characterizing at the transcriptional (RNA-seq) and phenotypic levels (estimation of autonomous seed production) a series of synthetic *Capsella* diploid and tetraploid hybrids. We found that the predicted dominance relationships matched the observed expression of self-incompatibility alleles, as well as the mating system phenotypes. Hence, the mating system of newly formed *Capsella* allotetraploids depends on the dominance relationship between self-incompatibility alleles inherited from the parents. Overall, our results improve our understanding of the mechanisms by which changes in ploidy can alter the system of mating over the course of evolution.

## Introduction

Mating systems have far-reaching effects on plant evolution (Wright et al. 2013). For instance, shifts from outcrossing to self-fertilization are expected to reduce the effective rate of recombination and genetic polymorphism (Glémin et al. 2006), while at the same time providing reproductive assurance when mates are scarce (Jain, 1976). The establishment of polyploid populations is an iconic example of these effects. Whole-genome duplication (WGD) is prevalent in plants (Soltis et al. 2015), and polyploid species are overrepresented in the Arctic flora (Brochmann et al. 2004) as well as in invasive (Pandit et al. 2011) and domesticated plants (Salman-Minkov et al. 2016). Moreover, ancient WGD events on phylogenies seem to be associated with drastic environmental changes (Vanneste et al. 2014). Therefore, WGD has often been hypothesized to allow faster adaptation and niche differentiation in changing environments (Selmecki et al.2015; Baniaga et al. 2020). However, beside these potential long-term advantages, newly formed polyploid genotypes are also expected to suffer from the immediate lack of gametes of the same cytotype and from the lower fitness of interploidy hybrids. This phenomenon, known as “minority cytotype exclusion” (Levin, 1975; Husband, 2000), is expected to drastically hinder the success of newly formed polyploid lineages.

Self-fertilization should greatly increase the establishment success of polyploid populations by allowing them to avoid minority cytotype exclusion (Fowler & Levin, 2016). However, empirical surveys on the association between polyploidy and self-fertilization either confirmed the positive association (Barringer, 2007; Robertson et al. 2011), or found no association (Mable, 2004). This suggests that the current models fail to incorporate important details of the interaction between polyploidy and self-fertilization, such as *e.g.* mechanisms controlling the mating system or the confounding effects of hybridization. An intriguing observation is that allopolyploid lineages (in which WGD occurred in association with hybridization) often exhibit low outcrossing rates, whereas autopolyploid lineages (where “only” WGD occurred) often exhibit predominant outcrossing or mixed mating systems (Husband et al. 2008).

The mating system of Brassicaceae species is controlled by a sporophytic self-incompatibility system, in which self-pollen recognition is caused by the allele-specific interaction between a pollen coat ligand protein (encoded by the *SCR* gene) and a stigma transmembrane receptor kinase (encoded by the *SRK* gene, Takayama et al. 2001). The two genes are tightly linked within a small genomic region called the S-locus, where a large number of S-alleles (also called S-haplotypes) typically segregate in self-incompatible species. S-haplotypes form a complex dominance hierarchy in anthers, whereby small non-coding RNA (sRNA) generated by dominant S-haplotypes transcriptionally silence the *SCR* gene of recessive S-haplotypes (Tarutani et al. 2010, Durand et al. 2014). Hence, while a large fraction of individuals are heterozygous at the S-locus, in most cases transcripts from only one of the two *SCR* alleles are present (Kakizaki et al. 2003; Burghgraeve et al. 2020), resulting in phenotypic dominance. Diversification of S-haplotypes in Brassicaceae is very ancient, as indicated by the very high level of nucleotide divergence among S-haplotype sequences and extensive trans-specific and even trans-generic sharing among related taxa (Castric & Vekemans, 2004). In *Arabidopsis* and *Capsella*, S-haplotypes are classified into four main dominance classes, related to their phylogenetic relationships with class I being the most recessive and class IV being the most dominant (Prigoda et al. 2005; Durand et al.2014; Bachmann et al. 2019). Several allopolyploid species of the Brassicaceae family originated from the hybridization between a self-incompatible (SI) and a self-compatible (SC) parental species, including *A. suecica* (with *A.thaliana* as SC parent and *A. arenosa* as SI parent; Novikova et al. 2017)*, A. kamchatica* (with *A. lyrata* as SC parent and *A. halleri* as SI parent; Shimizu-Inatsugi et al. 2009; Kolesnikova et al. 2022), and *C. bursa-pastoris* (with *C. orientalis* as SC parent and *C. grandiflora* as SI parent; Douglas et al. 2015). These three species have a recent allopolyploid origin, and all share the common feature of being self-compatible. The reason why these allopolyploid species originating from SIxSC are SC rather than SI is intriguing. An interesting possibility could be that the dominance interactions between S-haplotypes could have caused the instantaneous breakdown of SI in these species if the (non-functional) S-haplotype contributed by the SC species had retained the ability to suppress the expression of the (functional) *SCR* alleles contributed by the SI species in hybrid offspring (reviewed in Novikova et al. 2022). Consistent with this hypothesis, the allotetraploid *A. suecica, A. kamchatica,* and *C. bursa-pastoris* all share the same nonfunctional S-allele as their respective SC parental species (Novikova et al. 2017; Bachmann et al. 2019, 2021; Kolesnikova et al. 2022). In addition, at least some resynthesized *A. suecica*- or *C. bursa-pastoris*-like allotetraploids are immediately SC after hybridization (Novikova et al. 2017; Bachmann et al. 2021; Duan et al. 2023), also supporting that the loss of SI could be an instant outcome of possessing one (relatively dominant) non-functional S-haplotype. A key prediction from this scenario is that the mating system of the resulting hybrid should vary according to the dominance of the S-haplotype contributed by the SI parent relative to that of the SC parent. However, formal proof of this hypothetical process, *i.e.* the establishment of a direct causal link between the relative dominance of the S-haplotypes, the expression of *SCR* alleles in anthers, and the loss of SI in allopolyploids, is still lacking.

Here, we used an experimental approach based on a series of synthetic allopolyploid individuals obtained between the selfer *C. orientalis* and the outcrosser *C. grandiflora* (Duan et al. 2023) to test whether the breakdown of SI observed in allopolyploid species in Brassicaceae, such as *C. bursa-pastoris*, could be explained by the dominance interaction between S-haplotypes in anthers. First, we used published genomic and transcriptomic resequencing data to assess a methodology to infer S-locus genotypes and pollen S-locus phenotypes in *Capsella*. Then we used RNA-sequencing (RNA-seq) data from young inflorescences to measure allele-specific expression of the *SCR* and *SRK* genes in synthetic diploid and tetraploid *C. orientalis* x *C. grandiflora* hybrids, as an approximation of the early stages of natural *C. bursa-pastoris*. Finally, we compared the observed expression of *SCR* alleles with the seed set from autonomous self-pollination in the synthetic hybrids. Altogether, our results demonstrate that the relative dominance of S-alleles inherited from the parental species is a key determinant of the mating system of nascent polyploid lineages.

## Results

### A reliable methodology to determine S-locus genotypes and phenotypes in Capsella

First, we produced a comprehensive set of S-allele reference sequences in *Capsella*. For this, we genotyped individuals at the *SRK* gene using the NGSgenotyp pipeline (Genete et al. 2020) on publicly available short-read resequencing data of 180 *C. grandiflora* individuals from Monodendri, Greece (the Cg-9 population in Josephs et al. 2015), starting from a database of *SRK* sequences from *A. lyrata* and *A. halleri* and 62 partial *C. grandiflora SRK* sequences previously obtained by Sanger sequencing by Jesper Bechsgaard and Mikkel Schierup (Guo et al. 2009; Neuffer et al. 2023; see Supplementary Information). We obtained a fully resolved S-locus genotype for 177 individuals (Table S1), and identified 74 different S-alleles, including 25 previously unknown *C. grandiflora* S-alleles (Table S2). For most of them we were able to obtain full sequences of the exon 1 of *SRK* (available at https://www.doi.org/10.6084/m9.figshare.22567558). Among those new S-alleles we identified an apparently functional *SRK* allele, noted H4047, that shares 99% identity with the *SRK* pseudogene (for which the coding sequence is interrupted at position 949 of exon 1) found at the S-locus in subgenome A of *C. bursa-pastoris* by Bachmann et al. (2021). A more complete description of this set of S-alleles is given in the Supplementary Information. Beside these 74 *SRK* alleles, we also identified five sequences clustering with *SRK* alleles (H0002, H0003, H0011, H0012 and H0013 in Table S1, named CgrSRK01, CgrSRK06, CgrSRK09, CgrSRK51 and CgrSRK63 in Paetsch et al. 2006, and Neuffer et al. 2023, see Table S2) that we considered as paralogous sequences unlinked to the S-locus, as already documented in Schierup et al. (2001) and Prigoda et al. (2005, see Supplementary Information). Second, we validated the use of transcriptomic data to determine S-locus genotypes using the NGSgenotyp pipeline. We applied the approach described above to genotype the S-locus of four *C. grandiflora,* four *C. orientalis*, and 16 *C. bursa-pastoris* individuals, using published genome resequencing data as well as RNA-seq data obtained separately from leaf, root or flower bud tissues from the exact same individuals (Kryvokhyzha et al. 2019). Strictly identical S-locus genotypes were inferred based on genomic DNA and RNA-seq data from flower bud tissues (Table S3), as expected by the codominant expression of *SRK* in pistils reported in Brassicaceae (Hatakeyama et al. 2001; Burghgraeve et al. 2020), hence demonstrating that RNA-seq data from flower buds can be used to reliably genotype the S-locus in *Capsella*. We note that all four individuals of *C. grandiflora* were heterozygous at the S-locus, and all *C. orientalis* individuals were homozygous for the allele called H4004*n* (we use the notation “*n”* to indicate that this allele is non-functional), in agreement with Bachmann *et al*. (2019, 2021; note that these authors refer to this allele as CoS12). In agreement with Bachmann *et al*. (2021), most allotetraploid *C. bursa-pastoris* individuals had two copies of the H4004*n* allele derived from the non-functional *C. orientalis* parental allele and all individuals had two copies of the non-functional H4047*n* allele, derived from the functional *C. grandiflora* parental allele H4047 (Table S3). However, we found that a second allele, H2002, already known in *C. grandiflora*, appeared to be segregating at the S-locus in the *C. orientalis* subgenome (present in two copies in accession DUB-RUS9 and in one copy, together with H4004n, in LAB-RUS-4). As expected, we generally observed no or very low expression of *SRK* in leaves or roots (Table S3).

Third, we evaluated whether S-locus phenotypes in pollen could be assessed based on RNA-seq data. We obtained RNA-seq data from flower buds for seven diploid *C. grandiflora* individuals used as parents in the production of synthetic polyploids and hybrids with *C. orientalis* (see below), as well as for six synthetic autotetraploids (respectively Cg2 and Cg4 in Fig.1). We obtained the S-locus genotypes (at *SRK*) of these individuals using the NGSgenotyp pipeline, as described above. Five *C. grandiflora* S-alleles were found to segregate in this experimental material, again with approximately balanced transcript levels between both *SRK* alleles in heterozygotes (Fig.2, Table S4). Then, we obtained full *SCR* transcript sequences for each of the five S-alleles by applying the *de novo* assembly module of the NGSgenotyp pipeline (available at https://www.doi.org/10.6084/m9.figshare.22567558), based on a reference database of known *SCR* sequences from *A. halleri* and *A. lyrata* (see Supplementary Material). We used these new reference sequences to compare patterns of allele-specific expression for *SRK* and *SCR* (Table S4). In agreement with the results of Burghgraeve *et al*. (2020) in *A. halleri,* allele-specific expression was much more asymmetric between *SCR* alleles than between *SRK* alleles. Indeed, in all heterozygous individuals, one of the two alleles contributed over 99% of the total *SCR* transcript levels, with a single exception (individual Cg4-1-3), where the relative expression was still as high as 93% (Table S4). This very strong allelic imbalance was also found in tetraploid individuals, suggesting that a dominant allele is capable of repressing several co-occurring alleles at once. The identity of the predominantly expressed allele was fully concordant with expectations based on the predicted classes of dominance between S-alleles (Table S4). Even in cases where two S-alleles of the same dominance class co-occurred (e.g. H2008 and H2022 in individual Cg2-12-3, or H4015 and H4035 in Cg4-1-3), the asymmetry of the transcript levels remained very strong, hence enabling us to determine the dominance hierarchy among the five alleles, as follows: H4035 > H4015 > H2008 > H2022 > H1001. We also analyzed one diploid and one tetraploid *C. orientalis* individual (Table S4) and observed a single S-allele in pistil and pollen, H4004n, whose *SCR* sequence was fully identical to that reported by Bachmann *et al*. (2019). These results also confirm that both *SRK* and *SCR* are still expressed in *C. orientalis*, as reported by Bachmann *et al*. (2019), even though they are non-functional.

**Fig. 1.**
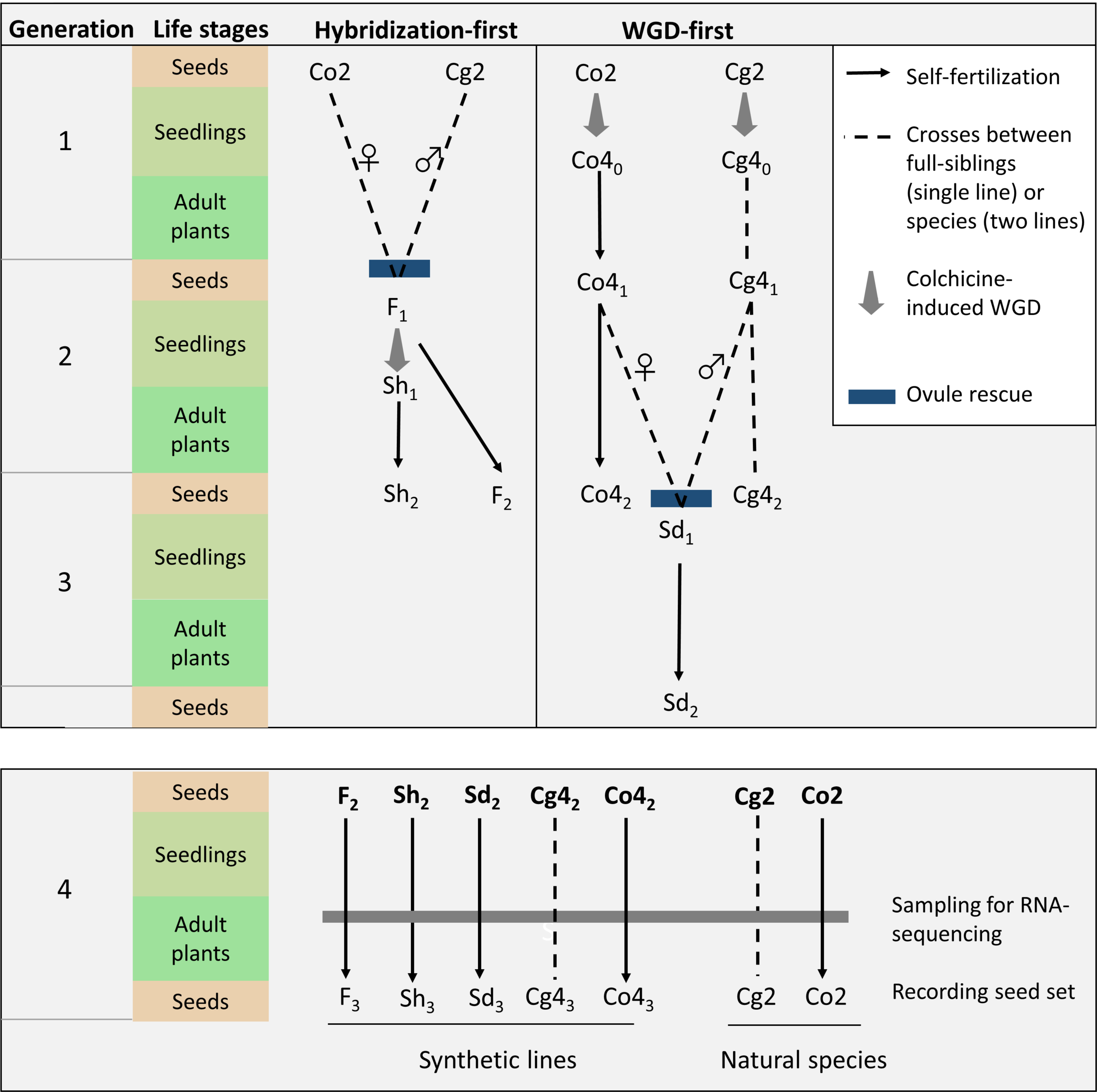
Crosses and induced whole-genome duplication (WGD) for generating diploid (F) and two types of tetraploid hybrids (Sd and Sh) from diploid *Capsella orientalis* (Co2) and *Capsella grandiflora* (Cg2; Duan et al. 2023). Autotetraploid *Capsella orientalis* (Co4) and *Capsella grandiflora* (Cg4) were created as intermediate steps. The generation numbers of the synthetic polyploids or hybrids are indicated by subscripts, and plant materials used in the present study are highlighted in bold.

**Fig. 2.**
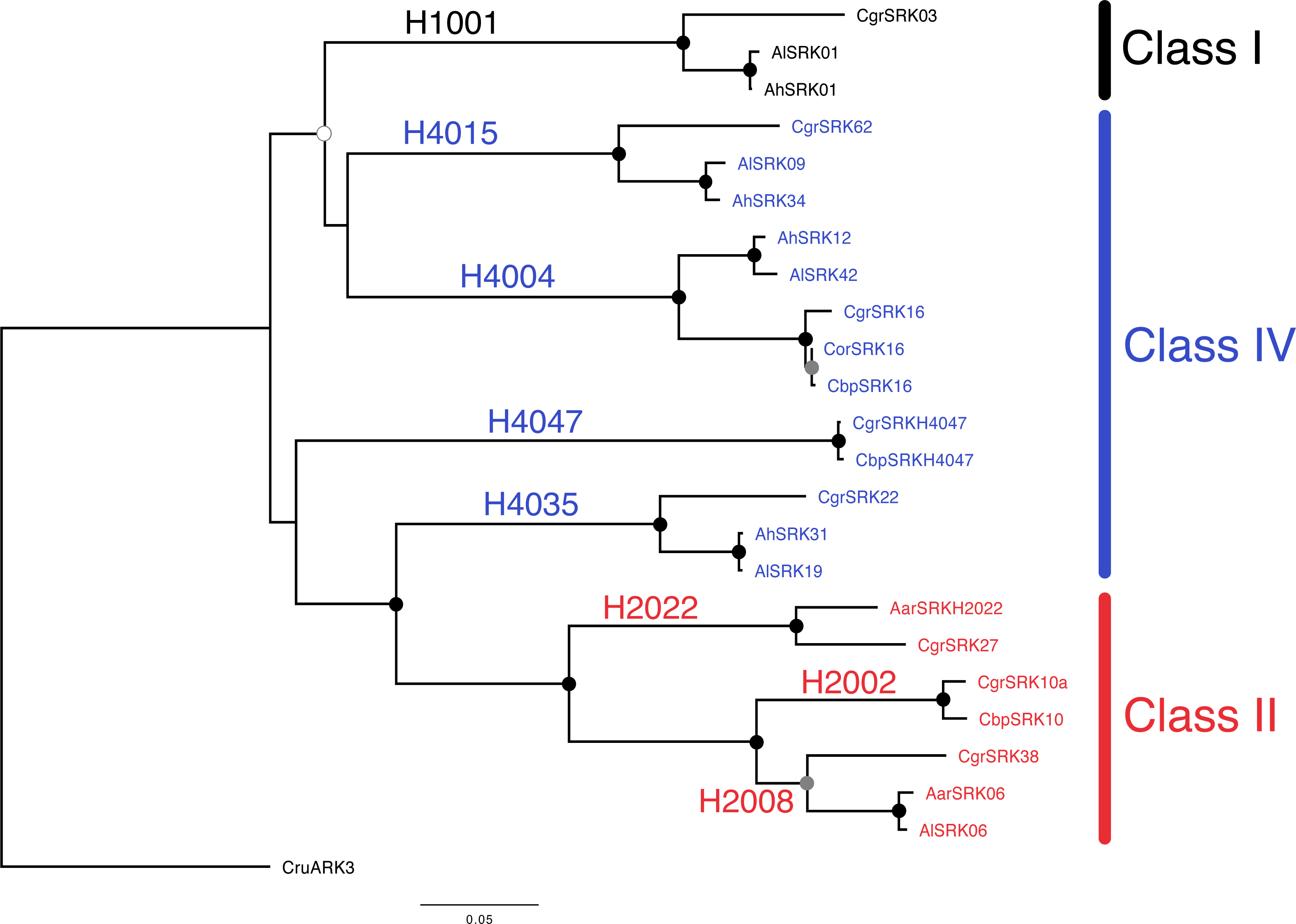
Phylogeny of *SRK* alleles identified in parents or hybrids between *Capsella grandiflora* (Cgr) and *C. orientalis* (Co). Phylogenetic reconstruction using maximum likelihood under the TPM3uf+I+G4 model implemented by PhyML (see Supplementary Information). We also included sequences identified in *C. bursa-pastoris* (Cbp), and trans-specific alleles from *Arabidopsis halleri* (Ah), *A. lyrata* (Al) or *A. arenosa* (Aar) when available. Functional specificities and dominance classes of the S-alleles are indicated. The outgroup is represented by an *ARK3* sequence from *Capsella rubella* (Cru). Bootstrap values are represented by dots on the nodes: black: BP = 100, grey: 95<BP<99, white: BP=89, no dot: BP < 80.

### The S-locus genotypes and phenotypes of diploid and synthetic tetraploid hybrids

Diploid and tetraploid hybrids were produced between *C. grandiflora* and *C. orientalis* (Fig. 1; Duan et al. 2023), using seeds from two wild individuals of *C. grandiflora* and one inbred line of *C. orientalis*. Diploid hybrids were generated by crossing *C. orientalis* with *C. grandiflora*, while allotetraploids were created either by inducing genome doubling in diploid hybrids with colchicine treatment (“Sh” allotetraploids), or by crossing colchicine-induced autotetraploid *C. orientalis* with autotetraploid *C. grandiflora* (“Sd” allotetraploids). In all crosses, diploid or tetraploid *C. orientalis* was used as the maternal plant, mimicking the formation of natural *C. bursa-pastoris* (Hurka et al. 2012). Then we analyzed RNA-seq data from flower buds of 27 diploid hybrids (F), seven tetraploidized hybrids (Sh) as well as 19 *C. orientalis* x *C. grandiflora* autotetraploid hybrids (Sd; Fig. 1, Tables 2, S5). Overall, five different S-alleles were segregating among these 53 individuals: the non-functional *C. orientalis* H4004*n* allele along with four *C. grandiflora* alleles (all alleles described above except H1001). Assuming that H4004n has retained the ability to transcriptionally repress S-alleles of a lower dominance class, as suggested by Bachmann *et al*. (2021), we used two different approaches to predict the SI phenotype of hybrids: (1) based on the S-locus genotype inferred from *SRK* data, a hybrid individual was predicted to be SC if it carried the *C. orientalis* H4004n allele and none of the S-alleles derived from *C. grandiflora* that are predicted to be more dominant than H4004n (i.e. H4035 and H4015), and to be SI otherwise (i.e. carrying H2008 and/or H2022); (2) based on the *SCR* relative expression data, a hybrid individual was predicted to be SC if the relative *SCR* read depth of allele H4004n was higher than 0.5, and to be SI otherwise. As shown in Table 2, predictions of the SI phenotypes based on the two methods were highly congruent. Overall, 18 of the 27 homoploid hybrids (F individuals), all 7 tetraploidized hybrids (Sh individuals), and 10 of the 19 hybrids between tetraploid *C. grandiflora* and *C. orientalis* (Sd individuals) were predicted to be functionally self-compatible. Indeed, in these individuals the presence of H4004n *SCR* transcripts was associated with efficient transcriptional repression of H2008 and H2022, as expected given their predicted classes of dominance. We observed only two exceptions to this general pattern (individuals F-5-6 and F-10-4, where H2022 had a higher level of expression than H4004n). Conversely, presence of H4015 and/or H4035 was associated with transcriptional repression of H4004n (Table 1), confirming recessivity of the latter with respect to these other class IV S-alleles.

**Table 1.** Prediction of mating system phenotype (self-compatible, SC; or self-incompatible, SI) of diploid and tetraploid hybrid individuals based on results on SRK genotype of individuals and previous knowledge about dominance relationships among S-alleles, and based on actual dominance of the non-functional allele copy of C. orientalis as estimated from relative expression levels of SCR. Grey boxes indicate the detection of the presence of an S-allele from SRK genotyping results. Numbers within boxes correspond to the mean read depth of SCR of the corresponding allele. Individual F-3-5 was suspected to be a spontaneous allotetraploid based on its phenotypes.

| Hybrid individual | Ploidy | Presence/absence of *SRK* and mean read depth of *SCR* allele: |  |  |  |  | *SCR* read depth of H4004n relative to total | Predicted SI/SC phenotype based on *SRK* genotype |
| --- | --- | --- | --- | --- | --- | --- | --- | --- |
|  |  | H2022 | H2008 | H4004n | H4015 | H4035 |  |  |
| F-1-3 | 2x | 2.4 |  | 23.4 |  |  | 0.905 | SC |
| F-1-4 | 2x | 0.0 |  | 1.8 |  |  | 1 | SC |
| F-1-5 | 2x |  |  | 105.6 |  |  | 1 | SC |
| F-1-6 | 2x | 0.0 |  | 3.6 |  |  | 1 | SC |
| F-3-1 | 2x |  |  | 10.2 |  |  | 1 | SC |
| F-3-5 | 2x | 3.0 |  | 59.9 |  |  | 0.953 | SC |
| F-5-1 | 2x | 0.0 |  | 18.1 |  |  | 1 | SC |
| F-5-3 | 2x | 0.0 | 1.2 | 32.3 |  |  | 0.965 | SC |
| F-5-4 | 2x | 0.0 | 194.0 |  |  |  | 0 | SI |
| F-5-5 | 2x | 1.2 | 468.5 |  |  |  | 0 | SI |
| F-5-6 | 2x | 141.4 |  | 21.2 |  |  | 0.13* | SC |
| F-8-2 | 2x |  |  |  | 1083.6 |  | 0 | SI |
| F-8-3 | 2x |  |  | 4.7 | 429.8 |  | 0.011 | SI |
| F-8-4 | 2x |  |  | 32.6 |  |  | 1 | SC |
| F-8-5 | 2x |  |  | 2.5 | 784.6 |  | 0.003 | SI |
| F-8-6 | 2x |  |  | 4.2 | 514.0 |  | 0.008 | SI |
| F-9-1 | 2x | 8.9 |  | 20.6 |  |  | 0.699 | SC |
| F-9-2 | 2x | 1.2 |  | 6.1 |  |  | 0.84 | SC |
| F-9-4 | 2x | 3.0 |  |  |  |  | 0 | SI |
| F-9-5 | 2x | 0.0 |  | 23.7 |  |  | 1 | SC |
| F-9-6 | 2x | 1.7 |  | 3.1 |  |  | 0.638 | SC |
| F-10-1 | 2x | 15.8 |  | 55.8 |  |  | 0.779 | SC |
| F-10-2 | 2x |  |  | 5.9 |  |  | 1 | SC |
| F-10-3 | 2x | 1242.3 |  |  |  |  | 0.000 | SI |
| F-10-4 | 2x | 36.8 |  | 25.1 |  |  | 0.405* | SC |
| F-10-5 | 2x |  |  | 63.5 |  |  | 1.000 | SC |
| F-10-6 | 2x | 2069.8 |  |  |  |  | 0.000 | SI |
| Sd-1-1 | 4x | 0.0 |  | 66.3 |  |  | 1.000 | SC |
| Sd-2-3 | 4x | 1.8 | 2.5 | 69.7 |  |  | 0.942 | SC |
| Sd-4-1 | 4x |  |  | 16.9 |  | 434.3 | 0.038 | SI |
| Sd-4-2 | 4x |  |  | 5.5 | 1541.3 |  | 0.004 | SI |
| Sd-4-3 | 4x |  | 1.8 | 31.0 |  | 326.2 | 0.086 | SI |
| Sd-4-5 | 4x |  |  | 5.4 | 19.1 | 467.8 | 0.011 | SI |
| Sd-4-6 | 4x |  |  | 1.8 | 505.4 |  | 0.004 | SI |
| Sd-6-1 | 4x |  | 0.0 | 64.3 |  |  | 1.000 | SC |
| Sd-6-3 | 4x |  | 0.0 | 22.6 |  |  | 1.000 | SC |
| Sd-6-4 | 4x |  | 0.0 | 32.3 |  | 471.9 | 0.064 | SI |
| Sd-6-5 | 4x |  | 0.0 | 9.7 |  |  | 1.000 | SC |
| Sd-6-6 | 4x |  |  | 9.7 |  | 285.4 | 0.033 | SI |
| Sd-7-1 | 4x |  | 0.0 | 6.7 |  | 297.7 | 0.022 | SI |
| Sd-7-3 | 4x |  | 0.0 | 44.0 |  |  | 1.000 | SC |
| Sd-7-5 | 4x |  |  | 11.9 |  | 595.1 | 0.020 | SI |
| Sd-8-1 | 4x |  | 0.0 | 34.9 |  |  | 1.000 | SC |
| Sd-8-4 | 4x | 0.0 |  | 61.1 |  |  | 1.000 | SC |
| Sd-8-5 | 4x | 0.0 | 0.0 | 68.6 |  |  | 1.000 | SC |
| Sd-8-6 | 4x | 1.2 |  | 124.9 |  |  | 0.991 | SC |
| Sh-1-2 | 4x | 1.2 |  | 64.9 |  |  | 0.982 | SC |
| Sh-2-5 | 4x | 0.6 |  | 37.3 |  |  | 0.984 | SC |
| Sh-3-5 | 4x | 2.4 |  | 39.0 |  |  | 0.942 | SC |
| Sh-5-5 | 4x | 0.0 | 0.0 | 141.9 |  |  | 1.000 | SC |
| Sh-7-5 | 4x | 0.0 |  | 99.4 |  |  | 1.000 | SC |
| Sh-7-6 | 4x | 1.1 |  | 45.6 |  |  | 0.976 | SC |
| Sh-9-2 | 4x | 0.0 | 0.0 | 31.9 |  |  | 1.000 | SC |

### Inferred S-locus phenotypes predict the mating system of hybrids

We then compared the autonomous seed set of diploid and tetraploid hybrids to test whether the *S*-locus genotypes and the observed transcriptional dominance of *SCR* alleles can explain which *C. orientalis* × *C. grandiflora* hybrids are SC and which are SI. Seed production under autonomous selfing was used as an indicator of self-compatibility and was classified into three categories to represent individuals with almost no seed (<10 seeds), few seeds (10-300 seeds) and a large number of seeds (> 300 seeds).

We found that both the *SRK*-predicted dominance of the H4004n allele and the relative expression level of its *SCR* allele are strong predictors of the ability of autonomous seed production, with only a few exceptions (Fig. 3). The *SRK*-based prediction of self-compatibility was strongly associated with seed production categories (Fisher’s exact test, p-value < 0.001), with most individuals predicted to be self-incompatible producing no or few seeds under autonomous pollination (<300 seeds). In contrast, individuals that were predicted to be self-compatible usually produced more than 300 seeds. Similarly, the relative expression level of the H4004n *SCR* allele significantly differed among the three seed production categories (Kruskal–Wallis H test, p-value < 0.001). Individuals with a higher proportion of the H4004n *SCR* allele expression (e.g., larger than 0.25) usually had autonomous seed production above 300, and individuals with a lower proportion of H4004n expression usually had no or few seeds.

**Fig. 3.**
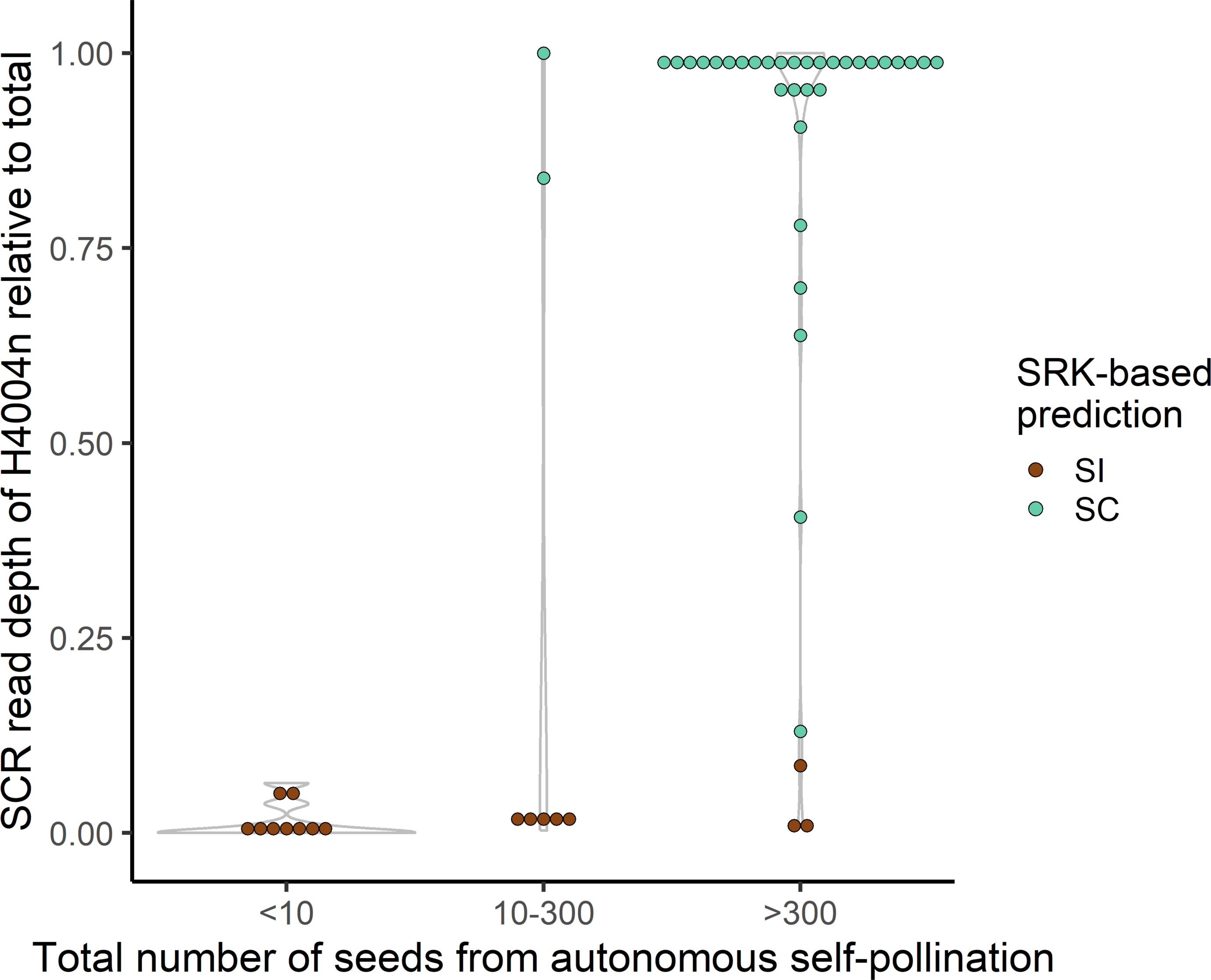
Relationships of the *SRK*-predicted dominance of the H4004n *SCR* allele, the observed relative expression level of the H4004n *SCR* allele and seed production in the diploid and tetraploid *C. orientalis C. grandiflora*.

## Discussion

### Assessing dominance relationships between S-alleles using RNA-seq data

At the S-locus in Brassicaceae, the genotype-to-phenotype map is complicated by the widespread existence of dominance/recessivity interactions between S-alleles. Determination of these dominance relationships ultimately relies on phenotypic assays based on controlled pollinations. Recently, Burghgraeve *et al*. (2020) demonstrated that phenotypic dominance in pollen can be predicted with high accuracy from the simple comparison of transcript abundances, using quantitative RT-PCR of *SCR* transcripts. However, allele-specific qPCR primers need to be designed and optimized for every single S-haplotype whose expression is to be quantified, which is not practical when large numbers of S-haplotypes segregate. Here, we show that RNA-seq data from flower buds can be used to reliably infer dominance relationships between S-alleles in pollen. These data are relatively simpler to obtain, as they do not require any specific optimization step beyond a generic RNA-seq library construction and can thus be generalized more readily than the qRT-PCR approach of Burghgraeve *et al*. (2020), or the labor-intensive phenotypic assessment of dominance. A limitation of this new approach, however, is that it can only quantify transcripts of *SCR* alleles whose nucleotide sequence is known *a priori*, which is only the case for a subset of the numerous S-haplotypes typically found in SI species, including *C. grandiflora*. A potential caveat to this new approach is that it relies on comparing mapping densities of (Illumina) sequencing reads on the nucleotide sequence of *SCR* alleles, which have relatively short coding sequences, thus making accurate mapping a potential challenge. The high levels of nucleotide divergence among *SCR* alleles is expected to (at least partially) compensate for this limitation, and accordingly, we found that for the five S-alleles considered in our crossing design, cross-mapping of individual sequencing reads among alleles is negligible, which makes the method highly reliable.

When applying the method to diploid *C. grandiflora* individuals, we found that the putatively dominant S-haplotype represented >99% of the global level of *SCR* transcripts in all individuals (Table S5), corresponding to nearly complete dominance at the transcriptional level, in line with Burghgraeve et al. (2020). The relative dominance levels we inferred among the *Capsella* S-haplotypes we studied here were also fully concordant with the dominance interactions previously measured by controlled crosses for the trans-specifically shared S-haplotypes in *A. lyrata* and *A. halleri* (Prigoda et al. 2005; Llaurens et al. 2008; Durand et al. 2014). Specifically, we confirmed that both S-haplotypes from class IV were dominant over both S-haplotypes from class II, which were themselves dominant over the single class I S-haplotype (Table S5). Another original result from our analysis is that a single *SCR* allele was majoritarily expressed in each tetraploid *C. grandiflora* individual, suggesting that the transcriptional silencing machinery controlling dominance remains effective in a tetraploid context, in line with the phenotypic patterns of dominance between S-haplotypes observed in tetraploid individuals of *A. lyrata* (Mable et al., 2004). We note that the exact number of gene copies of each allele observed in tetraploid individuals in our study remains uncertain because S-locus genotypes were determined using RNA-seq data instead of genomic resequencing (Genete et al. 2020). Hence, precise genotyping of tetraploid individuals based on genomic DNA will be needed to quantify the extent of this phenomenon.

The phenotypic effect of the observed dominance relationship among S-alleles was tested by measuring seed production in the diploid and tetraploid *C. orientalis* x *C. grandiflora* hybrids. Our measurement of seed production under autonomous selfing is not a formal test of SC, and several confounding effects could blur the link between the dominance relationship of *SCR* alleles and seed production. In particular, newly formed interspecific hybrids are expected to have lower fitness due to interactions between divergent genomes (Fishman & Sweigart, 2018), therefore individuals with no seed or few seeds could also result from hybrid incompatibility rather than SI. Furthermore, the hybrid individuals were not strictly separated in the growth chamber during flowering time, so for individuals that generated a small number of seeds, we cannot rule out the possibility of pollen contamination from other *Capsella* plants. Nonetheless, we still found a strong association between relative expression level of the non-functional H4004n *SCR* allele and autonomous seed production, which provides evidence that the machinery controlling dominance relationships between *SCR* alleles is a major determinant of the mating system of the hybrids we obtained.

### S-haplotypes dominance mediates the effect of WGD on the mating system of allopolyploids in the Brassicaceae

Associations between WGD and an autogamous mating system have been largely debated in plant biology, but a general consensus is still lacking (Mable, 2004; Barringer, 2007; Husband et al. 2008; Robertson et al. 2011). Part of the confusion stems from the fact that broad-scale studies focused on taxa comprising a mixture of different SI systems (e.g. self-recognition-based sporophytic SI, self-recognition or non-self recognition-based gametophytic SI), and different types of polyploidy (autopolyploidy, allopolyploidy, or a combination of both, i.e. segmental allopolyploidy). Moreover, the mechanistic causes of the loss of SI in polyploid lineages have generally remained elusive.

In the specific case of allopolyploidy in Brassicaceae, our results formally establish that the mating system immediately upon hybridization depends on the relative dominance of the non-functional allele inherited from the selfer *C. orientalis* as compared to that of the functional S-allele(s) inherited from the outcrosser *C. grandiflora*, in line with the model proposed by Novikova et al. (2022). This observation raises at least three intriguing questions. First, the non-functional S-allele needs to still retain the capacity to remain dominant. This is the consequence of the particular genetic architecture of dominance between S-haplotypes, where dominance modifiers (small non-coding RNAs) are decoupled from the gene they regulate (Billiard & Castric 2011). This particular genetic architecture of dominance might be less rare than it was long thought to be (Billiard et al. 2021). Second, the variation we observed relies on the fact that the non-functional S-haplotype that was fixed in *C. orientalis* has an intermediate level of dominance. If it had been the most recessive, then all allotetraploid individuals would by definition have inherited a more dominant S-haplotype from *C. grandiflora*, and would thus remain SI. In contrast, if *C. orientalis* had fixed the most dominant S-haplotypes, then all hybrids would have turned SC. This raises the question of why these SC species have fixed S-haplotypes at intermediate levels of dominance, while in SI species (including *C. grandiflora*) the most recessive S-allele typically has by far the highest frequency (Billiard et al. 2007). Some factors affecting the fixation probability of SC mutations have been studied by Tsuchimatsu & Shimizu (2013), but to the best of our knowledge, the effect of dominance on the fixation probability of SC mutations in a sporophytic SI system remains to be investigated formally. A third interesting question is why most known examples of recent allotetraploids in Brassicaceae involve hybridization between a selfer and an outcrosser (Novikova et al. 2022). A more general survey of recent allopolyploids would be needed to determine the generality of this pattern, but one tempting hypothesis is that, given an appropriate combination of S-haplotypes, this kind of crosses might instantaneously generate SC allotetraploid individuals avoiding minority cytotype exclusion (Novikova et al. 2022).

Determining whether differences in the intensity of genetic conflicts between the outcrosser and the selfer genomes (in particular over development of the endosperm, Rebernig et al. 2015) can oppose this selective advantage would be an interesting next step. Finally, a parallel can be drawn with the more general process of Haldane’s sieve, in which advantageous alleles tend to be fixed more readily in natural populations when they are dominant because they are directly exposed to natural selection (Haldane, 1924, 1927). Here in contrast, the selective advantage would go to hybrid lineages that have inherited from their SI parent a S-haplotype more recessive than the S-allele that was fixed in the selfing lineage.

A similar mechanism was proposed by Bachmann et al. (2021) for the evolution of selfing in the allotetraploid *C. bursa-pastoris* from *C. orientalis* and *C. grandiflora* parents. Intriguingly, while *C. bursa-pastoris* is a strong selfer, the S-haplotype it inherited from *C. grandiflora* (H4047) belongs to class IV, and thus would a priori be expected to be at least as dominant as the S-haplotype it inherited from *C. orientalis* (H4004n). Also, we identified one *C. bursa-pastoris* individual lacking the H4004n allele but carrying a (more recessive) class II allele (H2002, Table S3), presumably inherited from *C. orientalis*. Hence, other mechanisms might be needed to explain the loss of SI in the early development of the *C. bursa-pastoris* lineage. It should be noted, however, that dominance interactions between the class IV S-alleles have been firmly established at the phenotypic level for a small number of S-alleles only, so the possibility remains that some class IV S-alleles (in this particular case, H4047) could actually be more recessive than H4004n. This is suggested by the observation of a putative target of the small non-coding RNA produced by the *C. orientalis* S-haplotype in close proximity to the H4047n *SCR* pseudogene within the A subgenome of *C. bursa-pastoris* (Bachmann et al. 2021). As we found that both alleles (H4004 and H4047) are segregating in the Cg-9 population, it would be interesting to obtain living material carrying those alleles and perform controlled crosses to test their relative dominance.

## Material and methods

### Methodological approach to type S-alleles in Capsella experimental material based on RNA-seq data

To build an extended dataset of reference sequences of *SRK* from the self-incompatible species *Capsella grandiflora,* we genotyped 180 individuals of the Cg-9 population of *C. grandiflora* from Monodendri, Greece (Josephs et al. 2015) at the *SRK* gene with the NGSgenotyp pipeline (Genete et al. 2020) using raw Illumina reads available from Sequence Read Archive (SRA, Table S1). For the *SRK* reference database, we used available sequences of *SRK* from *A. lyrata* and *A. halleri* (Takou et al. 2021; Genete et al. 2020), and 62 partial sequences from *Capsella grandiflora* (Guo et al. 2009; Neuffer et al. 2023; see Table S2). Briefly, this pipeline filters raw reads with a dictionary of k-mers extracted from the reference database, then uses Bowtie2 to align filtered reads against each reference sequence from the database and produces summary statistics with Samtools (v1.4; Danecek et al. 2021) allowing it to identify S-alleles present in each individual. The pipeline NGSgenotyp also contains a *de novo* assembly approach module which produces full sequences of the S-domain of *SRK* for alleles present as partial sequences in the database as well as for newly identified S-alleles.

We then compared the results of the S-alleles genotyping approach obtained from either genomic DNA or RNA-seq data from flower buds, leaf and root tissues. For this we used published data from Kryvokhyzha et al. (2019) on four individuals each of *C. grandiflora* and *C. orientalis*, and on 16 individuals of *C. bursa-pastoris*. We applied the NGSgenotyp pipeline separately on each dataset, using the *SRK* reference database expanded with the *Capsella* S-allele sequences obtained above. Our analysis showed that S-allele typing based on RNA-seq data from flower buds gave identical results than those obtained from genomic DNA, so in the rest of the analyses we only used flower buds RNA-seq data.

### Creating and sequencing the transcriptome of synthetic hybrids and polyploids and assessing their mating system

To test the dominance relationship among SI alleles and its phenotypic consequences we used diploid and tetraploid hybrids of *C. orientalis* x *C. grandiflora*, using the synthetic hybrids generated by Duan *et al*. (2023; Fig. 1 and Table 2). We measured fruit set under autonomous selfing to test whether self-compatibility of these hybrids can be predicted by the dominance of *SCR* alleles, by comparing the *SCR* alleles identified in transcriptomes of inflorescences with fruit-set from spontaneous self-pollination in 27 diploid and 26 tetraploid hybrids.

In short, the diploid and tetraploid hybrids were generated from one inbred line of *C. orientalis* (URAL-RUS5), and seeds that were collected from two wild *C. grandiflora* individuals of the same population (85.1 and 85.24). Specifically, all the synthetic hybrids were descendants of three *C. grandiflora* individuals (85.1-5, 85.24-1, and 85.24-5), two of which (85.24-1 and 85.24-5) had the same maternal plant. Diploid hybrids (F) were obtained by crossing *C. orientalis* with *C. grandiflora*. Tetraploid hybrids (allotetraploids) were generated in two ways: in the first case the two diploid species were first crossed, and whole-genome duplication (WGD) was induced on the first generation of diploid hybrids with colchicine solution, resulting in “hybridization-first” synthetic allotetraploids (Sh); in the second case, WGD was induced in both diploid species, then the synthetic autotetraploids were crossed to obtain “WGD-first” allotetraploids (Sd). In addition, several diploid hybrids were suspected to have spontaneous WGD without colchicine treatment based on observations on organ size and the shape of trichomes, including individual F-3-5 which was used in the present study. In all interspecific crosses, diploid or tetraploid *C. orientalis* served as the maternal plant. The second generation of diploid hybrids, Sh-allotetraploids and Sd-allotetraploids as well as the diploid and tetraploid parental species were then grown together in a growth chamber. Each of the three hybrid groups was represented by six lines (independent hybridization events), and each line was represented by six individuals. The six individuals of the same line were full siblings from self-fertilization. An overview of the mating scheme used to create the different resynthesized hybrids and polyploids is given in Fig. 1.

The ability to generate seeds with only autonomous selfing was recorded for all hybrid individuals as a categorical factor. The hybrid individuals were classified into three categories based on the total number of seeds: 1) having almost no seed (< 10 seeds), 2) having few seeds (10-300 seeds), and 3) having plenty of seeds (> 300 seeds). As the hybrid individuals were not strictly separated in the growth chamber during flowering, the cutoff of ten seeds was applied to reduce noise from occasional pollen contamination from other *Capsella* plants. The seed set data of two individuals were removed from the dataset, because they were severely affected by disease during flowering time.

The first group of RNA-seq data was from Duan et al. (2023). Total RNA was extracted from the inflorescence of 6 diploid hybrids and 14 allotetraploids, using a cetyl-trimethyl-ammonium-bromide (CTAB) based method. Sequencing libraries were prepared with Illumina TruSeq Stranded mRNA (poly-A selection) kit, and sequenced on three NovaSeq 6000 S4 lanes with 150-bp paired-end reads (SNP&SEQ Technology Platform in Uppsala). One sequencing library was prepared and sequenced for each diploid sample, and two libraries were prepared for each tetraploid sample. On average 38.6 and 77.3 million read pairs were generated for the diploid and tetraploid samples, respectively.

To obtain a larger sample size, we performed a second group of RNA-seq on inflorescences of 33 additional hybrid individuals, including 20 diploid hybrids and 13 allotetraploids. Inflorescence samples of this second group were from the same experiment as those from the first group, and were collected at the same time, and stored at −80°C before sequencing. Total RNA was extracted using an RNeasy Plant Mini Kit (Qiagen). The library preparation and sequencing platform were the same as the first group, except that one library was prepared for each individual, regardless of ploidy level. Sequencing of the second group of inflorescence samples yielded an average library size of 97.4 million RNA-seq reads.

### Determination of S-locus genotype and phenotype in the Capsella hybrids and synthetic polyploids and confrontation with mating system phenotype assessments

In Brassicaceae, the self-incompatibility phenotype in pollen depends on complex dominance relationships among S-alleles, achieved through modifier genetic elements consisting in small RNAs encoded by precursors lying at the S-locus of dominant alleles and targeting the *SCR* gene of recessive alleles (Tarutani et al. 2010; Durand et al. 2014). Dominance in pollen is thus regulated at the transcriptional level, and is associated with very strong inhibition of mRNA production of recessive alleles (Burghgraeve et al. 2020), which could potentially be revealed by analyzing RNA-seq data. Hence, we tested this approach by performing S-allele typing with the NGSgenotyp pipeline using separately a *SRK* reference database, to determine the S-locus genotype of individuals (because *SRK* alleles are always co-expressed in the style, Hatakeyama et al. 2001), and a *SCR* reference database, to determine which S-allele is majoritarily expressed in pollen (and thus putatively dominant). We applied this approach to RNA-seq data from 8 diploid individuals (7 *C. grandiflora* + 1 *C. orientalis*) and 7 tetraploids (6 *C. grandiflora* + 1 *C. orientalis*) used as parents in the hybrid experiments (Duan et al. 2023), with the same *SRK* reference database as above enlarged with newly obtained *C. grandiflora* allele sequences, and for *SCR* with a reference database of sequences from *A. halleri* and *A. lyrata* (Genbank sequences). For *SCR*, because of the higher sequence divergence among S-alleles than for *SRK*, we modified the NGSgenotyp parameters by reducing the k-mer size used for filtering to a value of 15 (the default size used for *SRK* was 20). This allowed us, with the de novo assembly module of NGSgenotyp, to obtain full coding sequences of *SCR* for all *C. grandiflora* alleles present in the hybrids. In order to quantify the relative expression of *SCR* alleles, we mapped individual RNA-seq data against each reference *SCR* sequence with the genotyp module of NGSgenotyp and used the mean read depth delivered by Samtools (v1.4; Danecek et al. 2021) to compute the ratio of the mean read depth of the predominantly expressed allele (i.e., the putative dominant allele) to the sum of the read depths of all alleles present. We applied the same approach in synthetic tetraploid *C. grandiflora* individuals, but for *SRK* data we could only report the number and identity of alleles present, and thus it was not possible to precisely genotype individuals, i.e. to determine the number of gene copies of any given allele identified (when the total number of alleles detected in an individual was lower than 4, which was the case for all tetraploid individuals). Once the proposed approach was validated on *C. grandiflora* diploid and tetraploid individuals, we applied it to the experimental diploid and tetraploid hybrids. We identified two categories of hybrid individuals, in terms of pollen SI phenotype, depending on relative dominance levels of the inherited *C. grandiflora* and *C. orientalis* parental alleles: individuals with predominant expression of the *C. orientalis* allele, which is a non-functional S-allele; and individuals with predominant expression of one of the *C. grandiflora* alleles. Dominance of the non-functional *C. orientalis* allele is expected to cause a breakdown of self-incompatibility because it will impede recognition and rejection of self-pollen, and thus we assigned an expected self-compatible phenotype to those individuals, and an expected self-incompatible phenotype to individuals with a dominant *SCR* allele from *C. grandiflora*. Then we compared seed production data under autonomous selfing with the prediction of self-compatible/self-incompatible phenotype based on *SCR* dominance. The association between *SRK*-predicted self-compatibility and seed production categories was tested with Fisher’s exact test in R software environment version 3.6.3 (R Core Team, 2020). The relative expression level of the H4004n *SCR* allele (mean read depth of H4004n/sum of the mean read depth of all other S-alleles of the individual) in the three seed production categories were compared by the Kruskal–Wallis H test.

## Data availability

The RNA-sequencing data of the additional *Capsella* hybrids generated by this article are available in Sequence Read Archive (SRA) of National Center for Biotechnology Information (NCBI), and can be accessed with BioProject number PRJNA946929. All new Capsella *SRK* and *SCR* sequences obtained by de novo assembly with the NGSgenotyp pipeline are posted at https://www.doi.org/10.6084/m9.figshare.22567558.

## Author contributions

Conception and coordination of the study: TD, ML, VC, XV

Creation of experimental material: TD, AS

RNASEQ data: TD, ZZ

Bioinformatics: TD, MG

Data analysis: TD, XV, VC, CP, MG

Writing of the manuscript: TD, XV, VC, ML

All authors reviewed the manuscript.

## Acknowledgements

The work on SI in the Lille group is supported by the Agence Nationale de la Recherche (TE-MoMa project, grant number ANR-18-CE02-0020-01), the Région Hauts-de-France and the Ministère de l’Enseignement Supérieur et de la Recherche (CPER Climibio and CPER Ecrin grants), and the European Fund for Regional Economic Development. The study in Uppsala was supported by grant 2019-00806 from the Swedish Research Council to ML and Nilsson-Ehle research grant from the Royal Physiographic Society in Lund to TD. ML also acknowledges support from the Erik Philip-Sörensen Foundation. TD was supported by the Sven and Lilly Lawski Foundation. The computation and data handling were provided by the Swedish National Infrastructure for Computing (SNIC) at Uppmax, partially funded by the Swedish Research Council through grant agreement no 2018-05973. We also thank Elsa Sundkvist for her help with seed counting, Jesper Besgaard for early sharing of unpublished *SRK* sequences from *C. grandiflora*, and Tyler Kent and Stephen Wright for a preliminary analysis of S-alleles diversity in Cg-9 population. Finally, we thank Barbara Mable for her critical comments on the manuscript.

