## Supplementary Material for "Dominance between self-incompatibility alleles determines the mating system of Capsella allopolyploids"

### Supplementary information

#### Analysis of S-locus genotypes and quantification of relative transcript abundances using the bioinformatic pipeline NGSgenotyp

***Identification of S-alleles and classification into four dominance classes***

Despite the fact that most S-alleles are shared transpecifically between *A. halleri* and *A. lyrata* (Castric et al. 2008) and even transgenerically between *Arabidopsis* and *Capsella* (Paetsch et al. 2006), S-allele identifications/names have been assigned independently in each species, leading to much confusion in the literature. In this paper we introduce an identification scheme based on phylogenetic relationships among *SRK* sequences, in the form HXYYY, with X corresponding to the dominance class of the allele (ranging from 1 to 4), and YYY corresponding to its functional specificity (ranging from 001 up to a limit of 999). The functional specificity of *SRK* sequences from different species is assumed to be shared if they are phylogenetically closer to each other than to any other sequence from the same species (Castric et al. 2008). For instance, the most recessive allele in Arabidopsis and Capsella, called AhSRK01 in *A. halleri*, AlSRK01 in *A. lyrata* and CgrSRK03 in *C. grandiflora*, is here noted H1001 because it belongs to the dominance class I and is the first and only specificity in this class. The equivalence between species-specific identifications and our new generalized identification system is listed in Table S2. The dominance classes are also defined based on phylogenetic relationships among *SRK* sequences, as suggested initially by Prigoda et al. (2005) who defined four classes, as follows from the most recessive to the most dominant: A1, B, A3, A2. However, for more clarity and simplicity, we use instead the notation introduced by Goubet et al. (2012), from I to IV in the direction of increase in dominance: I (A1), II (B), III (A3) and IV (A2).

***Reference SRK and SCR sequences from Arabidopsis and Capsella***

Obtaining S-locus genotypes using the NGSgenotyp pipeline (Genete et al. 2020) requires a series of reference *SRK* and/or *SCR* sequences, so we started by assembling them from the literature. For *A. halleri*, the reference *SRK* sequences were taken from Table S1 of Genete et al. (2020). For *A. lyrata* they were taken from Table S14 of Mable et al. (2018) and from Takou et al. (2021). For *C. grandiflora* we used a set of partial S-domain sequences obtained by Paetsch et al. (2006), Guo et al. 2009, Bachmann et al. 2019, Nasrallah et al. 2007 and Neuffer et al. 2023. Reference sequences of *SCR* from *A. halleri* and *A. lyrata* were taken from Guo et al. (2011), Goubet et al. (2012), and Durand et al. (2014). All *SRK* and *SCR* reference sequences used are posted at https://www.doi.org/10.6084/m9.figshare.22567558 together with their Genbank accession numbers.

***S-locus genotypes and SRK sequences of the Cg-9 population of C. grandiflora***

We expanded the reference database of *Capsella* reference *SRK* sequences by using the *de novo* assembly module (haploasm) of the NGSgenotyp bioinformatic pipeline (Genete et al. 2020) based on Illumina short-read whole genome resequencing data of 180 individuals from the Cg-9 population of *C. grandiflora* from Monodendri, Greece (Josephs et al. 2015; Sequence Read Archive (SRA) BioProject PRJNA275635, Table S1). This allowed us to first obtain full sequences of exon 1 of *SRK* for the *C. grandiflora* for which only partial sequences were available (49 in total), as well as to obtain full sequences of exon 1 of *SRK* for the newly identified alleles (25 in total). The sequences of these 74 *SRK* alleles are available at https://www.doi.org/10.6084/m9.figshare.22567558. Based on this updated database of reference sequences, we then launched a new round of the genotyping module (genotyp) to obtain *SRK* genotypes. We obtained fully resolved diploid S-locus genotypes for 177 individuals (Table

S1), while sequence read depth was too low for two additional individuals. One individual was missing a single allele, and one individual had four alleles, which could correspond to an admixed sample or an autotetraploid individual.
Beside these 74 S-alleles, we also identified five sequences clustering with class II *SRK* alleles (H0002, H0003, H0011, H0012 and H0013 in Table S2, named, respectively, CgrSRK01, CgrSRK06, CgrSRK09, CgrSRK51 and CgrSRK63 in Paetsch et al. 2006 and Neuffer et al. 2023) that we considered as paralogous sequences unlinked to the S-locus for the following reasons : (1) two of them (H0002 and H0003) showed strong sequence similarity with sequences for which paralogy was confirmed with segregation analyses in *A. lyrata* and *A. halleri* (Aly13-2 and Aly13-7; Charlesworth et al., 2003; Castric & Vekemans, 2007); (2) the remaining three sequences had higher population frequencies than the other class II *SRK* alleles (11, 23 and 12 copies for H0011, H0012 and H0013, respectively, while true class II *SRK* alleles had frequencies ranging from 1 to 10 copies); and (3) they systematically occurred in individuals which already carried two other unambiguous *SRK* alleles (see Table S1).
Among the 74 S-alleles co-segregating in this population, one belongs to the most recessive class, 16 alleles belong to the second most recessive class (class II), 20 alleles belong to the more dominant class III and 37 alleles belong to the most dominant class IV (Table below). As theoretically expected (Schierup et al. 1997; Billiard et al. 2007), we observed that the more dominant classes contain a larger number of different S-alleles, each at a lower population frequency .

**Number and mean population frequency of S-alleles found in the population Cg-9 of *Capsella grandiflora* belonging to each of the four classes of dominance**

| | Class I | Class II | Class III | Class IV | Total |
|---|---|---|---|---|---|
| Number of S-alleles | 1 | 16 | 20 | 37 | 74 |
| Average frequency ± S.D. | 0.2436 | 0.0133 ± 0.0075 | 0.0115 ± 0.0065 | 0.0085 ± 0.0056 | / |

***Validation of the NGSgenotyp pipeline for use on RNA-seq data***
We evaluated how the NGSgenotyp performed when applied to RNA-seq data (from either flower bud, leaf or root tissues) by comparing the results with those obtained from whole genome resequencing data. We performed this comparison on four *C. grandiflora* and four *C. orientalis* individuals, as well as on 16 *C. bursa-pastoris* individuals from Kryvokhyzha et al. (2019, Table S1), for which both types of sequence data were available. We used as reference database of *SRK* sequences a combination of sequences from *A. halleri*, *A. lyrata*, and our enlarged set of *C. grandiflora* sequences.
For *C. grandiflora*, all four individuals were heterozygous at the S-locus, with allele H1001 shared by three individuals, while all other gene copies belonged to distinct S-alleles (H2002, H2006, H2017, H3014, and H4021) (Table S3). In addition, two individuals carried one copy each of putative paralogous sequences (H0011 and H0003 in individuals 85.3 and 86.12, respectively). For *C. orientalis*, all 4 individuals were homozygous for the non-functional allele H4004*n* (*n* is for non-functional), in agreement with Bachmann *et al.* (2019, note that these authors refer to this allele as CoS12). All *C. orientalis* individuals were also homozygous for an unlinked paralogous sequence (H0014, named CorSRK06 by Neuffer et al. 2023). Unlike the putative paralogous sequences observed in *C. grandiflora*, which are segregating at low or moderate frequencies at unlinked loci, the H0014 sequence appears to be fixed in *C. orientalis*. For the allotetraploid *C. bursa-pastoris*, in agreement with the results of Bachmann *et al.* (2021), all individuals but two had two copies of the H4004*n* allele, derived from the non-functional *C.*

*orientalis* parental allele, and all individuals had two copies of the H4047*n* allele, derived from the functional *C. grandiflora* parental allele H4047 (Table S3). Determining the exact copy number of alleles is possible using this dataset because analyses are based on genomic data, so that relative read depth informs about copy number (Genete et al. 2020). Indeed, as previously observed by Bachmann *et al.* (2021), accession DUB-RUS9 was lacking the H4004n allele inherited from *C. orientalis*, but we found that this individual was carrying two copies of a distinct allele, H2002, in the *C. orientalis* subgenome. Interestingly, another accession also carried one copy of the H2002 allele, LAB-RUS-4, but this accession also carried one copy of the *C. orientalis* H4004*n* and two copies of the *C. grandiflora* H4047 alleles. Hence, it appears that Russian accessions of *C. bursa-pastoris* have a second allele segregating at the S-locus of the *C. orientalis* subgenome, which was not reported by Bachmann et al. (2021). In addition, we found that all but five *C. bursa-pastoris* individuals have the *C. orientalis* subgenome fixed for the H0014 paralogous sequence, and all but two individuals were fixed, probably in the *C. grandiflora* subgenome, for the H0003 paralogous sequence (Table S3). The genomic locations of H0014 and H0003 were confirmed to be distinct from the *S*-locus region thanks to Blast analysis against a public genome assembly of *C. bursa-pastoris* (Kasianov et al. 2017).
Analysis of RNA-seq data from flower buds from the same set of individuals gave strictly identical results as those obtained from genomic data, confirming the codominant expression of *SRK* transcripts (Burghgraeve et al. 2020). As expected also, RNA-seq data detected no expression of *SRK* from leaf or root tissues, with only three exceptions: two S-alleles from class II (H2002 and H2006), and one S-allele from class III (H3014). The expression of *SRK* in leaf tissues was already reported for some *SRK* alleles in *A. lyrata*, all from class II (Prigoda et al. 2005). Overall, these results confirm that the NGSgenotyp pipeline can be used to reliably infer S-locus genotype from RNA-seq data from flower buds.

***Determining patterns of allelic expression in SRK and SCR in diploid and tetraploid parents.***
We combined RNA-seq data from seven diploid and six synthetic tetraploid individuals of *C. grandiflora* used as parents for the hybrid material with *C. orientalis* (Duan et al. 2023, BioProject PRJNA848625) with newly obtained RNA-seq data from additional hybrid individuals (BioProject PRJNA946929) and applied the NGSgenotyp pipeline to obtain their S-locus genotype by focusing first on *SRK*. Overall, five different S-alleles were segregating among the 13 individuals: H1001, H2008, H2022, H4015 and H4035 (Fig. 2). All these alleles were present in the Cg-9 population and thus their sequences were present in the database. We used the "average read depth" computed by the genotyping module of NGSgenotyp to quantify relative expression of the *SRK* alleles present within each diploid and tetraploid individual. The results showed overall co-dominant transcript levels of the different *SRK* alleles (Table S4). Note that in the tetraploid individuals analyzed, we observed up to only three different *SRK* alleles, which implies that one of the three is present in two copies. However, the use of RNA-seq data precludes reliable determination of *SRK* copy numbers, as can be done using genomic resequencing data (see above).
Then we aimed to obtain the associated *SCR* sequences, so applied again the *de novo* assembly module (haploasm) of NGSgenotyp on the same individuals, but this time using the library of *SCR* reference sequences from *A. halleri* and *A. lyrata* described above, and using shorter k-mers (length of 15 instead of the default value of 20). Although reference *SCR* sequences were available only for H1001 and H2008 (from *A. lyrata*), we successfully obtained full coding sequences for all five S-alleles. The sequences are available at https://www.doi.org/10.6084/m9.figshare.22567558. We used the *SCR* reference sequence database enriched with these five newly obtained *SCR* sequences to launch a new round of the genotyping module (genotyp) to estimate the relative expression of *SCR* alleles. The results showed strongly asymmetric allele-specific expressions (see main text and Table S4). The results also showed that only one allele was highly expressed in tetraploid individuals, suggesting that the small RNA-mediated regulation of *SCR* expression is fully functional in tetraploids. These results also allowed to determine the patterns of hierarchical dominance among these five S-alleles, as follows: (1) H4035 dominant over H4015 (individual cg4-1-3-F); (2) H4035 dominant over H2022 and H2008 (cg4-8-3-F, cg4-12-4-F, cg4-9-2-F); (3) H4015 dominant over H2022 and H2001 (cg2-1-2-F, cg2-2-6-F, cg2-1-6-F, cg2-7-3-F); (4)

H2008 dominant over H2022 (cg2-12-3-F, cg2-14-5-F, cg4-6-4-F, cg4-7-3-F). Hence, assuming transitivity of the dominance relationships: H4035 > H4015 > H2008 > H2022 > H1001.

***Determining patterns of allelic expression in SRK and SCR in diploid and tetraploid hybrids***
Finally, we applied the genotyp module of NGSgenotyp with *SRK* and *SCR* reference sequences, separately, on RNA-seq data from 27 diploid *C. orientalis* x *C. grandiflora* hybrids (F), seven tetraploidized hybrids (Sh) as well as 19 autotetraploid hybrids (Sd) obtained by crossing the parents described above (Fig. 1, Table 1). Overall, five different S-alleles were segregating among these hybrid individuals: the non-functional *C. orientalis* H4004n allele along with four *C. grandiflora* alleles, two of which that are predicted to be more recessive (H2008, H2022), and two of which that are predicted to be more dominant (H4015, H4035) than H4004n in pollen (Fig. 2). Among the 27 diploid hybrids (F individuals), 21 were carrying at least one copy of the H4004n allele, whereas six did not, so that the latter can be hypothesized to be SI. Accordingly, these six individuals expressed a functional *SCR* allele from *C. grandiflora*, either H2008, H2022 or H4015 (Table 1). Among the 21 homoploid hybrids carrying the H4004n allele, three individuals carried H4015, assumed to be dominant over H4004n. Hence, we can hypothesize that these three individuals are also SI, whereas the 18 remaining individuals are expected to be SC. Accordingly, these three individuals expressed the H4015 functional allele from *C. grandiflora*, thus confirming that H4015 is indeed dominant over H4004n. The remaining 18 individuals expressed predominantly the H4004n allele, with two exceptions: individuals F-5-6 and F-10-4 carried H2022 and H4004n but expression of the former was higher than that of the latter. All seven tetraploidized hybrids (Sh individuals) carried H4004n and the recessive H2022 allele, and are thus hypothesized to be SC. Again, *SCR* results confirm that these individuals expressed the H4004n allele only. Finally, the 19 hybrids between tetraploid individuals of *C. grandiflora* and *C. orientalis* (Sd individuals) were all found to carry H4004n, presumably inherited from their *C. orientalis* parent. Nine of them carried at least one of the more dominant *C. grandiflora* alleles H4015 or H4035, and can be assumed to be SI, whereas the other ten individuals received a more recessive S-allele from *C. grandiflora*, and are expected to be SC. These predictions were confirmed by *SCR* expression results (Table 1). Overall, in these hybrid individuals, the asymmetric expression of *SCR* sequences was strong, as observed above in *C. grandiflora* individuals, but the range of relative expression was wider (relative expression of the most dominant *SCR* allele ranging from 0.638 to 1.0) including in diploid hybrids, so this is not caused by polyploidy.

***S-haplotypes phylogeny based on full SRK exon1 sequences.***
In order to understand which S-haplotype specificities are shared within and among Arabidopsis and Capsella genera, 23 *SRK* (exon 1) nucleotide sequences from Arabidopsis and Capsella genera were aligned with Muscle v.3.5 (Edgar 2004) using the default strategy and manually checked with Seaview v.5 (Gouy et al. 2021). *ARK3* and *SRK* genes are paralogous therefore an *ARK3* sequence of *Capsella rubella* (Cru) was used as outgroup. On the alignment, Gblocks version 0.91b (Castresana 2000) was applied to remove poorly aligned regions. This resulted in a 1236 nucleotide dataset. The best fitting model under the ML criterion was selected with ModelTest-ng v.0.1.6 (Darriba et al. 2020). The phylogenetic reconstruction was performed by maximum likelihood (ML) using PhyML version 3.0 (Guindon et al. 2010) with the best of SPR and NNI moves on a BioNJ starting trees under the TPM3uf+I+G4 model. Node stability was estimated by 100 non-parametric bootstrap replicates.

**Cited in Supplementary Information**

Bachmann, J.A., Tedder, A., Fracassetti, M., Steige, K.A., Lafon-Placette, C., Köhler, C., et al. (2021) On the origin of the widespread self-compatible allotetraploid Capsella bursa-pastoris (Brassicaceae). Heredity, 127, 124–134.
Bachmann, J.A., Tedder, A., Laenen, B., Fracassetti, M., Désamoré, A., Lafon-Placette, C., et al. (2019) Genetic basis and timing of a major mating system shift in Capsella. New Phytologist, 224, 505–517.

Billiard, S., Castric, V. & Vekemans, X. (2007) A General Model to Explore Complex Dominance Patterns in Plant Sporophytic Self-Incompatibility Systems. Genetics, 175, 1351.
Charlesworth, D., Bartolome, C., Schierup, M.H. & Mable B.K. (2003) Haplotype Structure of the Stigmatic Self-Incompatibility Gene in Natural Populations of Arabidopsis lyrata. Molecular Biology and Evolution, 20, 1741–1753.
Castresana, J. (2000) Selection of conserved blocks from multiple alignments for their use in phylogenetic analysis. Molecular Biology and Evolution, 17, 540-552.
Castric, V., Bechsgaard, J., Schierup, M.H. & Vekemans, X. (2008) Repeated Adaptive Introgression at a Gene under Multiallelic Balancing Selection. PLOS Genetics, 4, e1000168.
Castric, V. & Vekemans, X. (2007) Evolution under strong balancing selection: How many codons determine specificity at the female self-incompatibility gene SRK in Brassicaceae? BMC Evolutionary Biology, 7, 1–15.
Darriba, D., Posada, P., Kozlov, A.M., Stamatakis, A., Morel, B. & Flouri, T. (2020) ModelTest-NG: A New and Scalable Tool for the Selection of DNA and Protein Evolutionary Models. Molecular Biology and Evolution, 37, 291–294.
Duan, T., Sicard, A., Glémin, S. & Lascoux, M. (2023) Expression pattern of resynthesized allotetraploid Capsella is determined by hybridization, not whole-genome duplication. New Phytologist, 237, 339–353.
Durand, E., Méheust, R., Soucaze, M., Goubet, P.M., Gallina, S., Poux, C., et al. (2014) Dominance hierarchy arising from the evolution of a complex small RNA regulatory network. Science, 346, 1200–1205.
Edgar, R.C. (2004) MUSCLE: Multiple Sequence Alignment with High Accuracy and High Throughput. Nucleic Acids Research, 32, 1792-7.
Genete, M., Castric, V. & Vekemans, X. (2020) Genotyping and De Novo Discovery of Allelic Variants at the Brassicaceae Self-Incompatibility Locus from Short-Read Sequencing Data. Molecular Biology and Evolution, 37, 1193–1201.
Goubet, P.M., Bergès, H., Bellec, A., Prat, E., Helmstetter, N., Mangenot, S., et al. (2012) Contrasted Patterns of Molecular Evolution in Dominant and Recessive Self-Incompatibility Haplotypes in Arabidopsis. PLOS Genetics, 8, e1002495.
Gouy, M., Tanier, E., Comte, N. & Parsons, D.P. (2021) Seaview Version 5: A Multiplatform Software for Multiple Sequence Alignment, Molecular Phylogenetic Analyses, and Tree Reconciliation in Methods in molecular biology (Clifton, N.J. editor) 2231, 241-260.
Guindon, S., Dufayard, J.F., Lefort, V., Anisimova, M., Hordijk, W. & Gascuel, O. (2010) New Algorithms and Methods to Estimate Maximum-Likelihood Phylogenies: Assessing the Performance of PhyML 3.0. Systematic Biology, 59, 307-321.
Guo, Y.L., Bechsgaard, J.S., Slotte, T., Neuffer, B., Lascoux, M., Weigel, D., et al. (2009) Recent speciation of Capsella rubella from Capsella grandiflora, associated with loss of self-incompatibility and an extreme bottleneck. Proceedings of the National Academy of Sciences of the United States of America, 106, 5246–5251.
Guo, Y.L., Zhao, X., Lanz, C. & Weigel, D. (2011) Evolution of the S-Locus Region in Arabidopsis Relatives. Plant Physiology, 157, 937–946.
Josephs, E.B., Lee, Y.W., Stinchcombe, J.R. & Wright, S.I. (2015) Association mapping reveals the role of purifying selection in the maintenance of genomic variation in gene expression. Proceedings of the National Academy of Sciences of the United States of America, 112, 15390–15395.
Kasianov, A.S., Klepikova, A.V., Kulakovskiy, I.V., Gerasimov, E.S., Fedotova, A.V., Besedina, E.G., et al. (2017) High-quality genome assembly of Capsella bursa-pastoris reveals asymmetry of regulatory elements at early stages of polyploid genome evolution. The Plant Journal, 91, 278–291.
Kryvokhyzha, D., Milesi, P., Duan, T., Orsucci, M., Wright, S.I., Glémin, S., et al. (2019) Towards the new normal: Transcriptomic convergence and genomic legacy of the two subgenomes of an allopolyploid weed (Capsella bursa-pastoris). PLoS Genetics, 15, e1008131.

Mable, B.K., Brysting, A.K., Jørgensen, M.H., Carbonell, A.K.Z., Kiefer, C., Ruiz-Duarte, P., et al. (2018) Adding complexity to complexity: Gene family evolution in polyploids. Frontiers in Ecology and Evolution, 6, 114.
Nasrallah, J.B., Liu, P., Sherman-Broyles, S., Schmidt, R. & Nasrallah, M.E. (2007) Epigenetic Mechanisms for Breakdown of Self-Incompatibility in Interspecific Hybrids. Genetics, 175, 1965.
Neuffer, B., Bechsgaard, J., Paetsch, M., Titel, C., Wesse, C., Bona, E., et al. (2023) S-alleles and mating system in natural populations of Capsella grandiflora (Brassicaceae) and its congeneric relatives. Flora, 299, 152206.
Paetsch, M., Mayland-Quellhorst, S. & Neuffer, B. (2006) Evolution of the self-incompatibility system in the Brassicaceae: identification of S-locus receptor kinase (SRK) in self-incompatible Capsella grandiflora. Heredity 2006 97:4, 97, 283–290.
Prigoda, N.L., Nassuth, A. & Mable, B.K. (2005) Phenotypic and Genotypic Expression of Self-incompatibility Haplotypes in Arabidopsis lyrata Suggests Unique Origin of Alleles in Different Dominance Classes. Molecular Biology and Evolution, 22, 1609–1620.
Schierup, M.H., Vekemans, X. & Christiansen, F.B. (1997) Evolutionary Dynamics of Sporophytic Self-Incompatibility Alleles in Plants. Genetics, 147, 835–846.
Takou, M., Hämälä, T., Koch, E.M., Steige, K.A., Dittberner, H., Yant, L., et al. (2021) Maintenance of Adaptive Dynamics and No Detectable Load in a Range-Edge Outcrossing Plant Population. Molecular Biology and Evolution, 38, 1820–1836.

### Supplemental Tables

**Table S1.** Fully resolved S-locus genotypes of 176 individuals from population Cg-9 of *Capsella grandiflora*, as obtained by the NGSgenotyp pipeline on whole-genome resequencing data from Josephs *et al.* (2015). Allele sequences are identified according to a Brassicaceae functional sequence group nomenclature, as determined based on sequence similarity with alleles from related species of *Capsella* and *Arabidopsis* genera (see Table S2 for correspondence with Capsella-specific S-allele IDs). The presence of putative polymorphic paralogous sequences (H0002=CgrSRK51, H0003=CgrSRK63, H0011=CgrSRK01, H0012=CgrSRK09, H0012=CgrSRK06) is also reported.

| Individual | SRA run | allele 1 | allele 2 | paralogous sequences |
|---|---|---|---|---|
| 1 | SRR2065265 | H2015 | H3001 | |
| 2 | SRR2070905 | H1001 | H4038 | |
| 3 | SRR2070909 | H1001 | H3001 | |
| 4 | SRR2065291 | H3001 | H2008 | H0002,H0011,H0012 |
| 5 | SRR2065298 | H1001 | H4026 | |
| 6 | SRR2070922 | H4030 | H4040 | |
| 7 | SRR2070923 | H1001 | H3029 | H0013 |
| 9 | SRR2065335 | H3014 | H2008 | |
| 10 | SRR2065181 | H1001 | H3007 | H0012 |
| 11 | SRR2065191 | H4010 | H4028 | H0011,H0012 |
| 13 | SRR2065208 | H4020 | H2021 | |
| 14 | SRR2065217 | H4010 | H3016 | |
| 15 | SRR2065226 | H4004 | H3005 | |
| 16 | SRR2065235 | H4019 | H4011 | |
| 17 | SRR2065247 | H3005 | H2021 | |
| 18 | SRR2065256 | H4020 | H1001 | |
| 19 | SRR2065263 | H3013 | H2017 | H0003 |
| 20 | SRR2065270 | H4001 | H4028 | |
| 23 | SRR2065271 | H3012 | H3022 | |
| 24 | SRR2065272 | H1001 | H2002 | H0012 |
| 25 | SRR2065273 | H1001 | H2021 | |

| | | | | |
|---|---|---|---|---|
| 26 | SRR2065274 | H2015 | H1001 | |
| 27 | SRR2065275 | H1001 | H4015 | H0013 |
| 28 | SRR2065276 | H4004 | H3002 | |
| 29 | SRR2065277 | H4008 | H3008 | H0012 |
| 30 | SRR2065278 | H3022 | H4006 | |
| 31 | SRR2065279 | H2006 | H4001 | |
| 32 | SRR2070910 | H3015 | H2017 | |
| 33 | SRR2065280 | H4024 | H3005 | H0012 |
| 35 | SRR2065282 | H1001 | H2008 | |
| 36 | SRR2065283 | H2010 | H2019 | |
| 37 | SRR2070911 | H3017 | H4045 | |
| 38 | SRR2065284 | H3023 | H2020 | |
| 39 | SRR2065285 | H2005 | H2010 | |
| 41 | SRR2065286 | H1001 | H4004 | |
| 42 | SRR2070912 | H2002 | H4017 | H0012 |
| 43 | SRR2065287 | H3013 | H4004 | H0012 |
| 44 | SRR2065288 | H1001 | H3005 | |
| 45 | SRR2070913 | H1001 | H4010 | |
| 46 | SRR2070914 | H1001 | H2017 | |
| 47 | SRR2070915 | H2002 | H4017 | |
| 47 | SRR2065289 | H1001 | H4002 | |
| 48 | SRR2070916 | H2011 | H4011 | |
| 49 | SRR2065290 | H3013 | H3002 | |
| 50 | SRR2065292 | H1001 | H1001 | |
| 51 | SRR2065293 | H1001 | H1001 | H0012 |
| 52 | SRR2065294 | H1001 | H1001 | |
| 53 | SRR2070918 | H1001 | H1001 | |
| 54 | SRR2065295 | H3029 | H3029 | H0012 |
| 55 | SRR2065296 | H4040 | H4046 | |
| 58 | SRR2065297 | H4027 | H3023 | |
| 59 | SRR2070919 | H1001 | H4040 | |
| 60 | SRR2070920 | H2010 | H4028 | |
| 61 | SRR2065299 | H1001 | H3016 | H0003 |
| 63 | SRR2065300 | H1001 | H2018 | |

| | | | | |
|---|---|---|---|---|
| 64 | SRR2065301 | H2017 | H2021 | |
| 65 | SRR2065303 | H1001 | H3029 | |
| 66 | SRR2065304 | H2012 | H3029 | |
| 67 | SRR2065305 | H1001 | H1001 | |
| 70 | SRR2065306 | H4040 | H3006 | |
| 71 | SRR2065307 | H1001 | H3006 | |
| 72 | SRR2065308 | H3005 | H4025 | |
| 74 | SRR2065309 | H3012 | H4037 | |
| 75 | SRR2065310 | H1001 | H2012 | |
| 76 | SRR2065311 | H1001 | H3015 | H0013 |
| 78 | SRR2065312 | H2015 | H3014 | |
| 79 | SRR2065313 | H2008 | H2018 | |
| 80 | SRR2065315 | H2002 | H3001 | |
| 81 | SRR2065316 | H1001 | H3001 | |
| 82 | SRR2065317 | H1001 | H4024 | |
| 83 | SRR2065318 | H1001 | H3005 | |
| 85 | SRR2065319 | H1001 | H3022 | |
| 86 | SRR2065320 | H2015 | H3005 | |
| 88 | SRR2070924 | H1001 | H4009 | |
| 89 | SRR2065321 | H2017 | H2021 | H0002 |
| 90 | SRR2065323 | H3015 | H3016 | |
| 91 | SRR2065324 | H4007 | H3023 | H0002 |
| 91 | SRR2070925 | H2006 | H4010 | |
| 92 | SRR2065325 | H3016 | H4038 | |
| 93 | SRR2065327 | H3017 | H3023 | |
| 94 | SRR2065328 | H1001 | H1001 | |
| 95 | SRR2065329 | H1001 | H2022 | |
| 96 | SRR2065330 | H1001 | H4028 | |
| 97 | SRR2065332 | H1001 | H3015 | H0002 |
| 98 | SRR2065333 | H4010 | H4028 | H0011 |
| 99 | SRR2065334 | H3005 | H4042 | H0011 |
| 101 | SRR2065172 | H2008 | H4030 | H0013 |
| 103 | SRR2065174 | H4001 | H4003 | H0003 |
| 105 | SRR2065175 | H4027 | H3029 | |

| | | | | |
|---|---|---|---|---|
| 106 | SRR2065176 | H1001 | H4035 | |
| 107 | SRR2065177 | H3017 | H4011 | |
| 108 | SRR2065179 | H4011 | H4030 | H0011, H0013 |
| 109 | SRR2065180 | H1001 | H1001 | |
| 110 | SRR2065182 | H1001 | H1001 | |
| 111 | SRR2065184 | H4010 | H4038 | H0002 |
| 112 | SRR2065185 | H1001 | H2017 | |
| 113 | SRR2065186 | H2002 | H4011 | |
| 115 | SRR2065188 | H4009 | H4047 | H0012 |
| 116 | SRR2070888 | H3015 | H4006 | |
| 117 | SRR2065189 | H1001 | H1001 | H0013 |
| 118 | SRR2065190 | H3005 | H4001 | |
| 119 | SRR2070889 | H2017 | H2010 | H0012 |
| 120 | SRR2070890 | H2010 | H2020 | H0003 |
| 121 | SRR2065192 | H2022 | H4046 | |
| 123 | SRR2065193 | H1001 | H2006 | |
| 124 | SRR2065195 | H4020 | H3006 | |
| 125 | SRR2065196 | H3002 | H4035 | H0013 |
| 126 | SRR2065197 | H1001 | H4028 | |
| 128 | SRR2065198 | H3010 | H4036 | |
| 129 | SRR2065199 | H4004 | H3007 | |
| 130 | SRR2070892 | H3001 | H2022 | |
| 131 | SRR2065200 | H2015 | H2020 | |
| 132 | SRR2065201 | H1001 | H3014 | |
| 133 | SRR2065202 | H1001 | H3003 | |
| 135 | SRR2065203 | H1001 | H4034 | |
| 136 | SRR2065204 | H3015 | H4047 | H0013 |
| 137 | SRR2065205 | H4017 | H2008 | H0013 |
| 138 | SRR2065206 | H4024 | H2020 | H0013 |
| 139 | SRR2065207 | H1001 | H3001 | H0002 |
| 140 | SRR2065209 | H3003 | H3013 | |
| 141 | SRR2065210 | H1001 | H4035 | H0012 |
| 142 | SRR2065211 | H1001 | H1001 | H0003,H0012, H0013 |
| 143 | SRR2065212 | H2011 | H2023 | |

| | | | | |
|---|---|---|---|---|
| 144 | SRR2065213 | H2012 | H4017 | |
| 145 | SRR2070893 | H4013 | H4030 | |
| 146 | SRR2065214 | H3001 | H2010 | |
| 147 | SRR2065215 | H3002 | H3029 | |
| 148 | SRR2065216 | H3010 | H2020 | |
| 149 | SRR2070894 | H1001 | H4037 | H0003 |
| 151 | SRR2065218 | H3016 | H4002 | |
| 152 | SRR2065219 | H4001 | H4045 | |
| 153 | SRR2065220 | H2010 | H4034 | |
| 154 | SRR2065221 | H1001 | H2022 | |
| 155 | SRR2065222 | H3003 | H3014 | |
| 156 | SRR2065223 | H4036 | H4003 | |
| 157 | SRR2065224 | H1001 | H4029 | |
| 158 | SRR2065225 | H4017 | H4006 | H0002 |
| 160 | SRR2065227 | H2002 | H4035 | H0012 |
| 161 | SRR2065228 | H1001 | H1001 | |
| 162 | SRR2065229 | H1001 | H3014 | H0003,H0011, H0012 |
| 163 | SRR2065230 | H1001 | H4014 | H0002 |
| 165 | SRR2065232 | H3011 | H3022 | |
| 166 | SRR2070895 | H4020 | H1001 | H0011, H0012 |
| 167 | SRR2065234 | H1001 | H2012 | |
| 168 | SRR2070896 | H2012 | H3019 | |
| 170 | SRR2065237 | H4027 | H4015 | |
| 172 | SRR2070897 | H3017 | H2018 | |
| 173 | SRR2065239 | H2012 | H4003 | H0002,H0012 |
| 174 | SRR2065241 | H4002 | H4013 | |
| 175 | SRR2065242 | H1001 | H4018 | H0012, H0013 |
| 176 | SRR2065244 | H2023 | H4028 | |
| 177 | SRR2065245 | H1001 | H2002 | |
| 178 | SRR2065246 | H4023 | H4029 | |
| 179 | SRR2070898 | H2010 | H2010 | H0012 |
| 180 | SRR2070900 | H1001 | H2020 | |
| 181 | SRR2065248 | H3003 | H4030 | H0003,H0012 |
| 182 | SRR2065250 | H1001 | H4039 | |

| | | | | |
|---|---|---|---|---|
| 183 | SRR2065251 | H4020 | H3029 | |
| 184 | SRR2065252 | H1001 | H4025 | |
| 186 | SRR2065254 | H1001 | H4001 | H0003 |
| 187 | SRR2065255 | H2013 | H4001 | |
| 189 | SRR2070901 | H1001 | H2022 | |
| 190 | SRR2065257 | H2015 | H2010 | |
| 192 | SRR2065258 | H1001 | H3011 | |
| 193 | SRR2065259 | H1001 | H4002 | |
| 195 | SRR2065260 | H2002 | H4001 | |
| 197 | SRR2065261 | H1001 | H3008 | |
| 198 | SRR2065262 | H1001 | H3003 | H0003 |
| 199 | SRR2070904 | H1001 | H2005 | |
| 200 | SRR2065266 | H1001 | H1001 | |
| 202 | SRR2065267 | H4001 | H2020 | H0012 |
| 203 | SRR2065268 | H4027 | H3010 | |
| 204 | SRR2065269 | H1001 | H4038 | |
| 207 | SRR2070906 | H2005 | H4026 | |
| 208 | SRR2070907 | H1001 | H4003 | |
| 209 | SRR2070908 | H4004 | H4003 | H0012 |

**Table S2.** List of the 74 *SRK* alleles of *C. grandiflora* identified within population Cg-9. References and Genbank accession numbers of previously published allele sequences are given, as well the source individual from population Cg-9 used to assemble a full sequence of exon 1 using the NGSgenotyp pipeline (available from file S1). References to five putative paralogous sequences are also given.

| Allele ID (functional group) | *Capsella* specific allele ID | Genbank | Original source | Individual used to extract full sequence | *A. halleri* homolog | *A. lyrata* homolog | Completeness of exon 1 sequence |
|---|---|---|---|---|---|---|---|
| H1001 | CgrSRK03 | DQ530639 | Paetsch et al. 2006 | 75_Contig_N1 | AhSRK01 | AlSRK01 | complete |
| H2002 | CgrSRK10 | MT592932 | Bechsgaard & Paetsch 2020 | 113_Contig_N5 | | | complete |
| H2005 | CgrSRK08 | MT592926 | Bechsgaard & Paetsch 2020 | 199_Contig_N7 | AhSRK23 | | complete |
| H2006 | CgrSRK24 | FJ649962 | Guo et al. 2009 | 123_Contig_N4 | AhSRK28 | AlSRK28 | complete |
| H2008 | CgrSRK38 | FJ649960 | Guo et al. 2009 | 101_Contig_N5 | | AlSRK06 | complete |
| H2010 | CgrSRK39 | MT592972 | Bechsgaard & Paetsch 2020 | 119_Contig_N4 | AhSRK69 | | complete |
| H2011 | CgrSRK34 | FJ613330 | Paetsch et al. 2006 | 143_Contig_N1 | | | complete |
| H2013 | CgrSRK23 | MT592948 | Bechsgaard & Paetsch 2020 | / | | | partial |
| H2015 | CgrSRK04 | DQ530640 | Paetsch et al. 2006 | 1_Contig_N3 | | | complete |
| H2017 | CgrSRK25 | MT592950 | Bechsgaard & Paetsch 2020 | 112_Contig_N3 | | | complete |
| H2018 | CgrSRK45 | MT592980 | Bechsgaard & Paetsch 2020 | 63_Contig_N5 | AhSRK68 | | complete |
| H2019 | CgrSRKH2019 | / | This study | 36_Contig_N3 | | | complete |
| H2020 | CgrSRK49 | MT592986 | Bechsgaard & Paetsch 2020 | 138_Contig_N5 | | | complete |
| H2021 | CgrSRK05 | DQ530641 | Paetsch et al. 2006 | 64_Contig_N3 | | | complete |
| H2022 | CgrSRK27 | MT592955 | Bechsgaard & Paetsch 2020 | 130_Contig_N6 | | | complete |
| H2023 | CgrSRKH2023 | / | This study | 143_Contig_N2 | | | complete |
| H2024 | CgrSRK40 | MT592976 | Bechsgaard & Paetsch 2020 | 75_Contig_N2 | | | complete |
| H3001 | CgrSRK18 | FJ649953 | Guo et al. 2009 | 130_Contig_N4 | AhSRK02 | AlSRK17 | complete |

| | | | | | | | |
|---|---|---|---|---|---|---|---|
| H3002 | CgrSRK17 | LR596609 | Bachmann et al. 2019 | / | AhSRK04 | AlSRK37 | complete |
| H3003 | CgrSRK12 | MT592934 | Bechsgaard & Paetsch 2020 | 140_Contig_N2 | AhSRK06 | AlSRK21 | complete |
| H3005 | CgrSRK32 | MT592960 | Bechsgaard & Paetsch 2020 | 86_Contig_N1 | AhSRK14 | | complete |
| H3006 | CgrSRKH3006 | / | This study | 70_Contig_N5 | AhSRK17 | | complete |
| H3007 | CgrSRKH3007 | / | This study | 10_Contig_N7 | AhSRK25 | | complete |
| H3008 | CgrSRKH3008 | / | This study | 29_Contig_N3 | AhSRK29 | AlSRK13 | complete |
| H3010 | CgrSRK46 | MT592982 | Bechsgaard & Paetsch 2020 | 203_Contig_N7 | AlSRK05 | | complete |
| H3011 | CgrSRK53 | MT592992 | Bechsgaard & Paetsch 2020 | 192_Contig_N6 | AhSRK57 | AlSRK25 | complete |
| H3012 | CgrSRK11 | FJ649956 | Guo et al. 2009 | 23_Contig_N4 | AhSRK60 | | complete |
| H3013 | CgrSRK13 | MT592935 | Bechsgaard & Paetsch 2020 | 140_Contig_N3 | | | complete |
| H3014 | CgrSRK19 | MT592944 | Bechsgaard & Paetsch 2020 | 132_Contig_N6 | AhSRK58 | | complete |
| H3015 | CgrSRK20 | MT592945 | Bechsgaard & Paetsch 2020 | 116_Contig_N4 | AhSRK70 | AlSRK70 | complete |
| H3016 | CgrSRK30 | MT592958 | Bechsgaard & Paetsch 2020 | 14_Contig_N1 | | | complete |
| H3017 | CgrSRK33 | MT592961 | Bechsgaard & Paetsch 2020 | 37_Contig_N1 | | | complete |
| H3019 | CgrSRK43 | MT592979 | Bechsgaard & Paetsch 2020 | 168_Contig_N4 | | | complete |
| H3022 | CgrSRKH3022 | / | This study | 165_Contig_N5 | AhSRK63 | AlSRK41 | complete |
| H3023 | CgrSRKH3023 | / | This study | 58_Contig_N5 | | AlSRK48 | complete |
| H3029 | CgrSRKH3029 | / | This study | 147_Contig_N6 | | | complete |
| H3030 | CgrSRK07 | EF530735 | Nasrallah et al. 2007 | 85_Contig_N2 | | | complete |
| H4001 | CgrSRK41 | MT592977 | Bechsgaard & Paetsch 2020 | 103_Contig_N2 | AhSRK05 | AlSRK34 | complete |
| H4002 | CgrSRK60 | MT593000 | Bechsgaard & Paetsch 2020 | 47_Contig_N2 | AhSRK07 | AlSRK72 | complete |
| H4003 | CgrSRKH4003 | / | This study | 103_Contig_N4 | AhSRK11 | AlSRK11 | complete |
| H4004 | CgrSRK16* | LR596551 | Bachmann et al. 2019 | / | AhSRK12 | AlSRK42 | complete |
| H4006 | CgrSRKH4006 | / | This study | 158_Contig_N6 | AhSRK15 | AlSRK71 | complete |
| H4007 | CgrSRK48 | MT592985 | Bechsgaard & Paetsch 2020 | 91_Contig_N5 | AhSRK16 | AlSRK31 | complete |
| H4008 | CgrSRK15 | MT592939 | Bechsgaard & Paetsch 2020 | 29_Contig_N4 | AhSRK18 | AlSRK39 | complete |
| H4009 | CgrSRK58 | MT592998 | Bechsgaard & Paetsch 2020 | 115_Contig_N4 | AhSRK20 | AlSRK69 | complete |

| | | | | | | | |
|---|---|---|---|---|---|---|---|
| H4010 | CgrSRK26 | MT592954 | Bechsgaard & Paetsch 2020 | 111_Contig_N3 | AhSRK21 | AlSRK15 | complete |
| H4011 | CgrSRK61 | MT593001 | Bechsgaard & Paetsch 2020 | 100_Contig_N5 | AhSRK22 | AlSRK46 | complete |
| H4012 | CgrSRK29 | MT592957 | Bechsgaard & Paetsch 2020 | 45_Contig_N7 | AhSRK24 | AlSRK20 | partial |
| H4013 | CgrSRKH4013 | / | This study | 174_Contig_N3 | AhSRK26 | AlSRK22 | complete |
| H4014 | CgrSRKH4014 | / | This study | 163_Contig_N5 | AhSRK32 | AlSRK68 | complete |
| H4015 | CgrSRK62 | MT593002 | Bechsgaard & Paetsch 2020 | 170_Contig_N2 | AhSRK34 | AlSRK09 | complete |
| H4017 | CgrSRKH4017 | / | This study | 42_Contig_N2 | AhSRK36 | AlSRK36 | complete |
| H4018 | CgrSRKH4018 | / | This study | 175_Contig_N4 | AhSRK37 | AlSRK27 | complete |
| H4019 | CgrSRK35 | FJ613331 | Paetsch et al. 2006 | 16_Contig_N3 | AhSRK38 | AlSRK65 | complete |
| H4020 | CgrSRK02 | DQ530638 | Paetsch et al. 2006 | 124_Contig_N7 | AhSRK39 | AlSRK35 | complete |
| H4023 | CgrSRK14 | FJ649958 | Guo et al. 2009 | 178_Contig_N2 | AhSRK43 | AlSRK43 | complete |
| H4024 | CgrSRK21 | MT592946 | Bechsgaard & Paetsch 2020 | 138_Contig_N4 | AhSRK47 | AlSRK63 | complete |
| H4025 | CgrSRKH4025 | / | This study | 72_Contig_N2 | AhSRK71 | AlSRK04 | complete |
| H4026 | CgrSRK31 | MT592959 | Bechsgaard & Paetsch 2020 | 5_Contig_N3 | AhSRK42 | AlSRK23 | complete |
| H4027 | CgrSRK37 | FJ649951 | Guo et al. 2009 | 58_Contig_N1` | AhSRK62 | AlSRK30 | complete |
| H4028 | CgrSRKH4028 | / | This study | 11_Contig_N2 | AhSRK51 | AlSRK33 | complete |
| H4029 | CgrSRKH4029 | / | This study | 157_Contig_N1 | AhSRK64 | AlSRK38 | complete |
| H4030 | CgrSRKH4030 | / | This study | 145_Contig_N6 | AhSRK50 | AlSRK50 | complete |
| H4034 | CgrSRK59 | MT592999 | Bechsgaard & Paetsch 2020 | 153_Contig_N6 | AhSRK54 | AlSRK73 | complete |
| H4035 | CgrSRK22 | MT592947 | Bechsgaard & Paetsch 2020 | 106_Contig_N7 | AhSRK31 | AlSRK19 | complete |
| H4036 | CgrSRK54 | MT592994 | Bechsgaard & Paetsch 2020 | 128_Contig_N3 | AhSRK55 | AlSRK12 | complete |
| H4037 | CgrSRKH4037 | / | This study | 74_Contig_N7 | AhSRK66 | AlSRK45 | complete |
| H4038 | CgrSRKH4038 | / | This study | 111_Contig_N4 | AhSRK52 | AlSRK44 | complete |
| H4039 | CgrSRK57 | MT592997 | Bechsgaard & Paetsch 2020 | 182_Contig_N4 | AhSRK56 | AlSRK61 | complete |
| H4040 | CgrSRKH4040 | / | This study | 6_Contig_N2 | AhSRK44 | AlSRK62 | complete |
| H4042 | CgrSRKH4042 | / | This study | 99_Contig_N5 | AhSRK46 | | complete |
| H4045 | CgrSRKH4045 | / | This study | 37_Contig_N5 | | | complete |
| H4046 | CgrSRKH4046 | / | This study | 121_Contig_N3 | | | complete |
| H4047 | CgrSRKH4047 | / | This study | 136_Contig_N7 | | | complete |

| | | | | | | |
|---|---|---|---|---|---|---|
| *putative paralogous sequences* | | | | | | |
| H0002 | CgrSRK51 | MT592989 | Bechsgaard & Paetsch 2020 | 139_Contig_N5 | Aly13-2 | complete |
| H0003 | CgrSRK63 | MW291556 | Neuffer et al. 2023 | 34_Contig_N3 | Aly13-7 | complete |
| H0011 | CgrSRK01 | DQ530637 | Paetsch et al. 2006 | 11_Contig_N1 | | complete |
| H0012 | CgrSRK09 | MT592927 | Bechsgaard & Paetsch 2020 | 119_Contig_N2 | | complete |
| H0013 | CgrSRK06 | DQ530642 | Paetsch et al. 2006 | 101_Contig_N3 | | complete |

*CgSRK16 has been named CgS12 by Bachmann *et al.* (2019) in analogy to its closely trans-specific sequence AhSRK12 from *A. halleri*.

**Table S3.** S-locus genotypes of individuals from *C. grandiflora*, *C. orientalis* and *C. bursa-pastoris*, as inferred from genomic DNA or RNA-seq data from flower buds, leaf or root tissues (based on data from Kryvokhyzha et al. 2019). References and specific notations for *C. grandiflora* alleles are given in Table S2. H4004n is a non-functional allelle present in *C. orientalis* and *C. bursa-pastoris* (Bachmann et al. 2019; Bachmann et al. 2021), derived from the functional H4004 allele from *C. grandiflora*. H4047n is a non-functional allele present in *C. bursa-pastoris*, that has not yet been identified in *C. grandiflora* (Bachmann et al. 2021). The presence of paralogous sequences (H0003, H0011, H0014) is also indicated (references given in Table S2; H0014 is equivalent to CorSRK06 described in Neuffer et al. 2022).

| **Species (Geographical group*)** | **Accession ID** | **Genomic DNA** | | **RNA flower buds** | | **RNA leaf** | | **RNA root** | |
|---|---|---|---|---|---|---|---|---|---|
| | | allele 1 *(paralogues)* | allele 2 (allele 3) | allele 1 | allele 2 (allele 3) | allele 1 | allele 2 | allele 1 | allele 2 |
| *C. grandiflora* | 85.3 | H2002 *(H0011)* | H4021 | H2002 *(H0011)* | H4021 | H2002 *(H0011)* | / | H2002 *(H0011)* | / |
| *C. grandiflora* | 86.12 | H1001 *(H0003)* | H2006 | H1001 *(H0003)* | H2006 | H2006 | / | H2006 | / |
| *C. grandiflora* | 87.26 | H1001 | H3014 | H1001 | H3014 | H3014 | / | H3014 | / |
| *C. grandiflora* | 88.5 | H1001 | H2017 | H1001 | H2017 | / | / | / | / |
| *C. orientalis* | GUB-RUS5 | H4004n *(H0014)* | H4004n | H4004n *(H0014)* | H4004n | / *(H0014)* | / | / *(H0014)* | / |
| *C. orientalis* | PAR-RUS | H4004n *(H0014)* | H4004n | H4004n *(H0014)* | H4004n | / | / | / | / |

| Species (Geographical group*) | Accession ID | Genomic DNA | | RNA flower buds | | RNA leaf | | RNA root | |
|---|---|---|---|---|---|---|---|---|---|
| *C. orientalis* | QH-CHIN4 | H4004n *(H0014)* | H4004n | H4004n *(H0014)* | H4004n | / *(H0014)* | / | / | / |
| *C. orientalis* | URAL-RUS4 | H4004n *(H0014)* | H4004n | H4004n *(H0014)* | H4004n | / | / | / | / |
| *C. bursa-pastoris* (ASI) | DL174 | H4047n *(H0003)* | H4004n *(H0014)* | H4047n *(H0003)* | H4004n *(H0014)* | / | / | / | / |
| *C. bursa-pastoris* (ASI) | JZH152 | H4047n *(H0003)* | H4004n *(H0014)* | H4047n *(H0003)* | H4004n *(H0014)* | / | / | / | / |
| *C. bursa-pastoris* (ASI) | NJ219 | H4047n *(H0003)* | H4004n *(H0014)* | H4047n *(H0003)* | H4004n *(H0014)* | / | / | / | / |
| *C. bursa-pastoris* (ASI) | TY118 | H4047n *(H0003)* | H4004n *(H0014)* | H4047n *(H0003)* | H4004n *(H0014)* | / | / | / | / |
| *C. bursa-pastoris* (CASI) | DUB-RUS9 | H4047n *(H0003)* | H2002 *(H0014)* | H4047n *(H0003)* | H2002 *(H0014)* | / | / | H2002 | / |
| *C. bursa-pastoris* (CASI) | KYRG-3-14 | H4047n | H4004n *(H0014)* | H4047n | H4004n *(H0014)* | / | / | / | / |
| *C. bursa-pastoris* (CASI) | LAB-RUS-4 | H4047n | H4004n (H2002) | H4047n *(H0003)* | H4004n (H2002) | / | / | H2002 | / |
| *C. bursa-pastoris* (CASI) | TACH-CHIN14 | H4047n *(H0003)* | H4004n | H4047n *(H0003)* | H4004n | / | / | / | / |
| *C. bursa-pastoris* (EUR) | FR50 | H4047n *(H0003)* | H4004n | H4047n *(H0003)* | H4004n | / | / | / | / |
| *C. bursa-pastoris* (EUR) | SE33 | H4047n *(H0003)* | H4004n | H4047n *(H0003)* | H4004n | / | / | / | / |

| Species (Geographical group*) | Accession ID | Genomic DNA | | RNA flower buds | | RNA leaf | | RNA root | |
|---|---|---|---|---|---|---|---|---|---|
| *C. bursa-pastoris* (EUR) | STA4 | H4047n *(H0003)* | H4004n | H4047n *(H0003)* | H4004n | / | / | / | / |
| *C. bursa-pastoris* (EUR) | STJ2 | H4047n *(H0003)* | H4004n *(H0014)* | H4047n *(H0003)* | H4004n *(H0014)* | / | / | / | / |
| *C. bursa-pastoris* (ME) | AL87 | H4047n *(H0003)* | H4004n *(H0014)* | H4047n *(H0003)* | H4004n *(H0014)* | / | / | / | / |
| *C. bursa-pastoris* (ME) | JO56 | H4047n *(H0003)* | H4004n *(H0014)* | H4047n *(H0003)* | H4004n *(H0014)* | / | / | / | / |
| *C. bursa-pastoris* (ME) | JO59 | H4047n *(H0003)* | H4004n *(H0014)* | H4047n *(H0003)* | H4004n *(H0014)* | / | / | / | / |
| *C. bursa-pastoris* (ME) | TR73 | H4047n *(H0003)* | H4004n *(H0014)* | H4047n *(H0003)* | H4004n *(H0014)* | / | / | / | / |

*Geographical groups of *C. bursa-pastoris* : ASI, Asian; CASI, Central Asion; EUR, European; ME, Middle East

**Table S4.** Relative expression levels of co-existing S-alleles in *SRK* and *SCR* in diploid and tetraploid individuals of *C. grandiflora* (cg2 and cg4) and *C. orientalis* (co2 and co4). The predominantly expressed allele in pollen is compared to putative dominance determined by previous knowledge about the dominance classes of the alleles. The difference in expression of the dominant allele is reported as a ratio of mean read depth, obtained by the genotyping pipeline NGSgenotyp.

| Individual | ploidy | allele 1 | *SRK* read depth | *SCR* read depth | allele 2 | *SRK* read depth | *SCR* read depth | allele 3 | *SRK* read depth | *SCR* read depth | Putative dominant allele in pollen | Allele expressed predomi-nantly in pollen | Relative SCR expression level of the dominant allele |
|---|---|---|---|---|---|---|---|---|---|---|---|---|---|
| cg2-1-2-F | 2x | H4015 | 15.12 | 931.38 | H2022 | 76.60 | 0.00 | | | | H4015 | H4015 | 1.0000 |
| cg2-1-6-F | 2x | H4015 | 13.06 | 608.58 | H1001 | 3.39 | 0.62 | | | | H4015 | H4015 | 0.9990 |
| cg2-2-6-F | 2x | H4015 | 7.53 | 671.57 | H2022 | 50.05 | 0.00 | | | | H4015 | H4015 | 1.0000 |
| cg2-7-3-F | 2x | H4015 | 22.58 | 745.59 | H1001 | 7.29 | 0.00 | | | | H4015 | H4015 | 1.0000 |
| cg2-9-5-F | 2x | H2022 | 94.62 | 5.19 | H1001 | 5.79 | 0.00 | | | | H2022 | H2022 | 1.0000 |
| cg2-12-3-F | 2x | H2008 | 6.47 | 710.67 | H2022 | 35.86 | 0.61 | | | | H2008 or H2022* | H2008 | 0.9991 |
| cg2-14-5-F | 2x | H2008 | 1.91 | 643.73 | H2022 | 90.21 | 5.30 | | | | H2008 or H2022* | H2008 | 0.9918 |
| cg4-1-3-F | 4x | H4035 | 20.55 | 520.44 | H4015 | 10.06 | 12.41 | H2022 | 22.28 | 3.77 | H4035 or H4015* | H4035 | 0.9375 |
| cg4-6-4-F | 4x | H2008 | 4.68 | 755.40 | H2022 | 49.90 | 3.92 | | | | H2008 or H2022* | H2008 | 0.9948 |

| | | | | | | | | | | | | | |
|---|---|---|---|---|---|---|---|---|---|---|---|---|---|
| cg4-7-3-F | 4x | H2008 | 7.60 | 2020.56 | H2022 | 21.03 | 0.60 | | | | H2008 or H2022* | H2008 | 0.9997 |
| cg4-8-3-F | 4x | H4035 | 35.39 | 1846.76 | H2022 | 12.85 | 0.00 | H2008 | 6.41 | 1.22 | H4035 | H4035 | 0.9965 |
| cg4-12-4-F | 4x | H4035 | 7.47 | 481.21 | H2022 | 9.56 | 0.00 | H2008 | 4.45 | 0.61 | H4035 | H4035 | 0.9908 |
| cg4-9-2-F | 4x | H4035 | 45.29 | 971.34 | H2008 | 15.60 | 0.00 | | | | H4035 | H4035 | 1.0000 |
| co2-2-6-F | 2x | H4004n | 20.56 | 32.21 | | | | | | | H4004n | H4004n | 1.0000 |
| co4-1-4-F | 4x | H4004n | 12.63 | 25.82 | | | | | | | H4004n | H4004n | 1.0000 |

*When two alleles belonging to the same dominance class co-occur, we cannot determine a priori which one is the most dominant, but *SCR* expression data allows us to resolve this uncertainty.

**Table S5.** S-locus genotypes and inference of putative self-compatible (SC) or self-incompatible (SI) phenotype of diploid and tetraploid individuals from *C. grandiflora* (*cg2* and *cg4*), *C. orientalis* (*co2* and *co4*) and experimental hybrids (F, Sd, Sh), as inferred from RNA-seq data from flower buds and analysis of *SRK* with the NGSgenotyp pipeline. The putative self-incompatible (SI) vs self-compatible (SC) phenotype of a given individual is inferred from previous knowledge of relative dominance of S-alleles in the pollen and from whether the non-functional allele H4004n from *C. orientalis* is present and putatively dominant in this individual. The presence of paralogous sequences (H0002, H0003, H0011, H0012, H0013, H0014) is also indicated.

| Individual | ploidy | allele 1 | allele 2 | allele 3 | allele 4 | putative paralogous sequences | Putative dominant allele | Putative phenotype |
|---|---|---|---|---|---|---|---|---|
| cg2-1-2-F | 2x | H4015 | H2022 | | | H0011,H0013 | H4015 | SI |
| cg2-1-6-F | 2x | H4015 | H1001 | | | H0011 | H4015 | SI |
| cg2-2-6-F | 2x | H4015 | H2022 | | | H0002,H0011 | H4015 | SI |
| cg2-7-3-F | 2x | H4015 | H1001 | | | H0002,H0011 | H4015 | SI |
| cg2-9-5-F | 2x | H2022 | H1001 | | | H0002,H0012 | H2022 | SI |
| cg2-12-3-F | 2x | H2008 | H2022 | | | H0011 | H2008 or H2022 | SI |
| cg2-14-5-F | 2x | H2022 | | | | H0011,H0012 | H2022 | SI |
| cg4-1-3-Fa | 4x | H4035 | H4015 | H2022 | | H0011,H0013 | H4035 or H4015 | SI |
| cg4-6-4-Fa | 4x | H2022 | H2008 | | | H0011 | H2008 or H2022 | SI |
| cg4-7-3-Fa | 4x | H2022 | H2008 | | | H0011 | H2008 or H2022 | SI |
| cg4-8-3-Fa | 4x | H4035 | H2022 | H2008 | | H0011,H0012 | H4035 | SI |
| cg4-12-4-Fa | 4x | H4035 | H2022 | H2008 | | H0011 | H4035 | SI |
| cg4-9-2-Fa | 4x | H4035 | H2008 | | | H0002,H0011 | H4035 | SI |
| co2-11-1-F | 2x | H4004n | | | | H00014 | H4004n | SC |
| co2-2-6-F | 2x | H4004n | | | | H00014 | H4004n | SC |
| co2-4-3-F | 2x | H4004n | | | | H00014 | H4004n | SC |
| co2-6-4-F | 2x | H4004n | | | | H00014 | H4004n | SC |
| co2-8-5-F | 2x | H4004n | | | | H00014 | H4004n | SC |

| | | | | | | |
|---|---|---|---|---|---|---|
| co2-9-5-F | 2x | H4004n | | H00014 | H4004n | SC |
| co4-1-4-F | 4x | H4004n | | H00014 | H4004n | SC |
| co4-3-1-F | 4x | H4004n | | H00014 | H4004n | SC |
| co4-4-6-F | 4x | H4004n | | H00014 | H4004n | SC |
| co4-8-3-F | 4x | H4004n | | H00014 | H4004n | SC |
| co4-9-3-F | 4x | H4004n | | H00014 | H4004n | SC |
| F-1-3 | 2x | H4004n | H2022 | H0002,H0012,H0014 | H4004n | SC |
| F-1-4 | 2x | H4004n | H2022 | H0002,H0012,H0014 | H4004n | SC |
| F-1-5 | 2x | H4004n | | H0012 | H4004n | SC |
| F-1-6 | 2x | H4004n | H2022 | H0002,H0012 | H4004n | SC |
| F-3-1 | 2x | H4004n | | H0012,H0014 | H4004n | SC |
| F-3-5 | 2x | H4004n | H2022 | H0012,H0014 | H4004n | SC |
| F-5-1 | 2x | H4004n | H2022 | H0011,H0014 | H4004n | SC |
| F-5-3 | 2x | H4004n | H2022 | H0011,H0014 | H4004n | SC |
| F-5-4 | 2x | H2022 | | H0011,H0014 | H2022 | SI |
| F-5-5 | 2x | H2008 | H2022 | H0011,H0014 | H2008 or H2022 | SI |
| F-5-6 | 2x | H4004n | H2022 | H0014 | H4004n | SC |
| F-8-2 | 2x | H4015 | | H0012,H0014 | H4015 | SI |
| F-8-3 | 2x | H4004n | H4015 | H0012,H0014 | H4015 | SI |
| F-8-4 | 2x | H4004n | | H0012,H0014 | H4004n | SC |
| F-8-5 | 2x | H4004n | H4015 | H0012,H0014 | H4015 | SI |
| F-8-6 | 2x | H4004n | H4015 | H0012 | H4015 | SI |
| F-9-1 | 2x | H4004n | H2022 | H0002,H0012 | H4004n | SC |
| F-9-2 | 2x | H4004n | H2022 | H0002,H0012 | H4004n | SC |
| F-9-4 | 2x | H2022 | | H0002,H0014 | H2022 | SI |
| F-9-5 | 2x | H4004n | H2022 | H0002,H0014 | H4004n | SC |
| F-9-6 | 2x | H4004n | H2022 | H0002,H0012,H0014 | H4004n | SC |
| F-10-1 | 2x | H4004n | H2022 | | H4004n | SC |
| F-10-2 | 2x | H4004n | | H0014 | H4004n | SC |
| F-10-3 | 2x | H2022 | | H0014 | H2022 | SI |

| | | | | | | | |
|---|---|---|---|---|---|---|---|
| F-10-4 | 2x | H4004n | H2022 | | H0014 | H4004n | SC |
| F-10-5 | 2x | H4004n | | | H0014 | H4004n | SC |
| F-10-6 | 2x | H2022 | | | H0014 | H2022 | SI |
| Sd-1-1 | 4x | H4004n | H2022 | | H0011,H0014 | H4004n | SC |
| Sd-2-3 | 4x | H4004n | H2022 | H2008 | H0011 | H4004n | SC |
| Sd-4-1 | 4x | H4004n | H4035 | | H0012,H0014 | H4035 | SI |
| Sd-4-2 | 4x | H4004n | H4015 | | H0011,H0012,H0014 | H4015 | SI |
| Sd-4-3 | 4x | H4004n | H4035 | | H0011,H0014 | H4035 | SI |
| Sd-4-5 | 4x | H4004n | H4035 | H4015 | H0012,H0014 | H4035 or H4015 | SI |
| Sd-4-6 | 4x | H4004n | H4015 | | H0011,H0012,H0014 | H4015 | SI |
| Sd-6-1 | 4x | H4004n | H2008 | | H0014 | H4004n | SC |
| Sd-6-3 | 4x | H4004n | H2008 | | H0014 | H4004n | SC |
| Sd-6-4 | 4x | H4004n | H4035 | H2008 | H0014 | H4035 | SI |
| Sd-6-5 | 4x | H4004n | H2008 | | H0014 | H4004n | SC |
| Sd-6-6 | 4x | H4004n | H4035 | | H0014 | H4035 | SI |
| Sd-7-1 | 4x | H4004n | H4035 | H2008 | H0011,H0014 | H4035 | SI |
| Sd-7-3 | 4x | H4004n | H2008 | | H0011,H0014 | H4004n | SC |
| Sd-7-5 | 4x | H4004n | H4035 | | H0014 | H4035 | SI |
| Sd-8-1 | 4x | H4004n | H2008 | | H0002,H0011,H0014 | H4004n | SC |
| Sd-8-4 | 4x | H4004n | H2022 | | H0002,H0012,H0014 | H4004n | SC |
| Sd-8-5 | 4x | H4004n | H2022 | H2008 | H0002,H0014 | H4004n | SC |
| Sd-8-6 | 4x | H4004n | H2022 | | H0002,H0014 | H4004n | SC |
| Sh-1-2 | 4x | H4004n | H2022 | | H0012,H0014 | H4004n | SC |
| Sh-2-5 | 4x | H4004n | H2022 | | H0002,H0011,H0014 | H4004n | SC |
| Sh-3-5 | 4x | H4004n | H2022 | | H0012,H0014 | H4004n | SC |
| Sh-5-5 | 4x | H4004n | H2022 | H2008 | H0011,H0014 | H4004n | SC |
| Sh-7-5 | 4x | H4004n | H2022 | | H0011 | H4004n | SC |
| Sh-7-6 | 4x | H4004n | H2022 | | H0011,H0014 | H4004n | SC |
| Sh-9-2 | 4x | H4004n | H2022 | H2008 | H0011,H0012,H0014 | H4004n | SC |
